# Impacts of different *SETD1A* mutations and genetic backgrounds on human induced pluripotent stem cell-derived hippocampal neurons

**DOI:** 10.64898/2026.10.07.756381

**Authors:** Yan Hong, Ting Zhao, Qian Yang, Hanwen Zhang, Xueting Ma, Xiao Su, Jiayi Liu, Siwei Zhang, Karla Mendez-Maldonado, Yi Zhou, Jizhong Zou, Anat Kreimer, Akira Sawa, Kimberly M. Christian, Zhiping P. Pang, Jubao Duan, Hongjun Song, Guo-li Ming

## Abstract

Human induced pluripotent stem cells (hiPSCs) provide a translational platform to interrogate the impact of genetic risk factors for various disorders on human biology. It has been a common practice to engineer premature stop codons to model predicted loss-of-function mutations using isogenic hiPSCs, yet effects of mutation position for the same gene and genetic background have been rarely tested systematically and at multiple levels for comparison to those of patient-specific mutations. Ultra-rare, heterozygous, and predicted loss-of-function mutations in the *SETD1A* gene, which encodes a H3K4 methyltransferase, are highly penetrant genetic risk factors for schizophrenia. Here, we generated a large panel of isogenic heterozygous *SETD1A* mutant hiPSC lines, including premature stop-codon mutations in one early exon and one late exon in five genetic backgrounds and two patient-specific mutations, to gain insight into molecular, cellular, and functional changes in a disease-relevant and highly pure population of human hippocampal dentate granule cell-like neurons. Our multi-level analyses showed that different *SETD1A* mutations lead to divergent molecular and cellular phenotypes related to synapses, but convergent neuronal network deficits, while the genetic background influences gene expression with no substantial effects at cellular and network levels. Mechanistic analysis confirmed the early exon patient mutation as loss-of-function, but revealed that the predicted loss-of-function patient mutation in the late exon is likely gain-of-function. Consequently, while all mutations decreased global H3K4me3 levels, deficits were only partially rescued by inhibiting H3K4 demethylase LSD1 for early exon mutations, but not for late exon mutations. Together, these results suggest that highly penetrant mutations can lead to similar cellular and functional phenotypes across different genetic backgrounds, whereas different premature stop-codon mutations could have contrasting impacts due to distinct mechanisms. Our findings have practical implications for applying genome-editing in hiPSCs to model disorders, highlight the value of modeling patient-specific mutations and testing multiple mutations, and suggest the need for tailored mutation-dependent therapeutic strategies.

## Main

The technology of human induced pluripotent stem cells (hiPSCs) reprogrammed from somatic cells, including from patients, to provide a genetically defined and renewable source of cells, has revolutionized our ability to study the biology of human cell types with limited accessibility, as well as to investigate biological mechanisms and treatment strategies for human diseases in cells with the same genetic composition as patients^1–4^. With a large number of genetic risk factors for many disorders recently identified, understanding how these genetic risk factors impact disease-relevant cell types not only provides insight into the etiology and pathophysiology of these disorders, but also may lead to novel therapeutic targets and treatment strategies. While patient-derived hiPSC lines can be generated for phenotypic analyses and drug screens, one major challenge has been to gain access to samples of patients with highly penetrant but very rare genetic mutations for hiPSC generation. The advancement of genome editing technologies, such as CRISPR/Cas9, has enabled engineering of specific mutations in hiPSCs to model these rare and highly penetrant mutations with isogenic lines to test their impact in many different genetic backgrounds^5–8^. Given that many patient mutations are predicted to be loss-of-function mutations, it has been a common practice to engineer premature stop codons to model these patient mutations^9–13^. Whether premature stop mutations in different positions lead to different outcomes and whether they indeed model patient-specific mutations have been rarely tested with multiple levels of analyses, especially in multiple genetic backgrounds.

Schizophrenia (SCZ), a highly heritable disorder with a ∼1% worldwide prevalence^14^, is widely considered a neurodevelopmental disorder arising from deficits in early brain development^15–17^. While many genetic risk factors for SCZ have been identified, our understanding of their biological functions lags behind^18–21^. A large-scale exome sequencing analysis from 24,248 SCZ cases and 97,322 controls identified rare coding variants in 10 genes that confer substantial risk for SCZ, with *SETD1A* as the top gene^22^. This study corroborates early genetic findings of the association of heterozygous premature stop mutations of *SETD1A*, which are predicted to be loss-of-function, with significant SCZ risk in smaller cohorts^21, 23^. *SETD1A* encodes a component of a histone methyltransferase complex that produces mono-, di-, and trimethylated histone 3 at lysine 4 (H3K4), a mark that on a genome-wide scale is broadly associated with transcriptional regulation and epigenetic tagging of promoter and enhancer sequences, and is proposed to perturb the expression of many target genes when disrupted^24–27^.

A number of rodent models with heterozygous deletions of different exons of *Setd1a* (*Setd1a^+/-^*) revealed several SCZ-related phenotypes, including working memory deficits^28^, impaired sociality^29^ and sensorimotor gating^30^. The common finding of a morphological defect in prefrontal cortical neurons, including decreased spine density, suggests that *Setd1a* haploinsufficiency results in synaptic deficits. Four previous studies have modeled *SETD1A* haploinsufficiency in hiPSCs. One study generated a heterozygous frameshift mutation in exon 7 of *SETD1A* by CRISPR/Cas9 in one hiPSC line and one human embryonic stem cell line, resulting in premature stop codons and *SETD1A* haploinsufficiency^12^. *SETD1A*-haploinsufficiency neurons dissociated from cortical spheroids showed reduced neurite outgrowth and decreased spontaneous activity by calcium imaging. The second study used CRISPR/Cas9 to generate a heterozygous frameshift mutation in exon 7 of *SETD1A* in two hiPSC lines from neurotypical subjects, also leading to a premature stop codon and *SETD1A* haploinsufficiency^6^. They generated composite networks of human induced glutamatergic and GABAergic neurons by forced expression of the transcription factor NGN2 or ASCL1, respectively. Transcriptomic analysis from one mutant line revealed dysregulated synaptic genes and an association of differentially expressed genes (DEGs) with SCZ. This mutant line showed enhanced dendritic complexity and elevated bursting activity by multielectrode array (MEA). A third recent study engineered a recurrent, patient-specific splice-site exon 16 mutation (c.4582-2delAG) into hiPSCs from a neurotypical subject. Differentiating hiPSCs into 2D neurons and 3D cerebral organoids showed a mitotic delay in neural progenitors accompanied by accelerated neurogenesis. The mutant cortical networks displayed robust hyperexcitability via MEA, but demonstrated reduced neurite complexity that contrasted with previous induced-neuron models^31^. A fourth recent study engineered the same patient-specific splice-site exon 16 mutation (c.4582-2delAG) and a heterozygous frameshift exon 16 mutation (p.L1533fs) in *SETD1A,* which did not alter the density of synapses but led to reduced dendritic complexity, increased firing rates and altered short-term synaptic plasticity revealed by patch-clamp recording^32^. Together, these early hiPSC studies revealed the complexity of *SETD1A* mutations on neuronal phenotypes, with some apparently contrasting findings. Because SCZ risk variants are found in multiple exons of *SETD1A*^21, 23^, whether introduction of premature stop mutations in different positions in the *SETD1A* gene leads to differential outcomes and whether the genetic background is a major determinant for phenotypes remain unknown.

To interrogate the impact and mechanism of *SETD1A* mutations, a relatively homogeneous and disease-relevant cell type is ideal for detailed cellular, molecular, epigenetic, and functional analyses. The hippocampus is implicated in the pathophysiology of SCZ, and hippocampal glutamatergic neurons, including dentate granule cells (DGCs), are functionally and structurally affected in SCZ^33–36^. Here, we developed a robust protocol for differentiating hiPSCs into hippocampal DGC-like neurons. Using CRISPR/Cas9 engineering, we generated 9 isogeneic pairs of hiPSC lines harboring heterozygous frameshift mutations targeting either early exon 4 or late exon 16 to generate premature stop codons in 5 different genetic backgrounds, including four neurotypical subjects and one idiopathic SCZ patient. We also generated 2 patient-specific mutations^21^ in exon 2 or exon 16 in one genetic background. We subsequently conducted a comprehensive investigation of the impact of different *SETD1A* mutations in DGC-like neurons at molecular, cellular, and functional levels. Our study illustrates that different *SETD1A* mutations result in divergent molecular and cellular phenotypes associated with synapses, but convergent neuronal network functional deficits, whereas the genetic background influences gene expression with no substantial effects at cellular and neuronal network levels. Our mechanistic study further uncovered that an exon 2 patient mutation represents loss-of-function, whereas an exon 16 patient mutation, predicted as a loss-of-function mutation, is likely gain-of-function. Our results highlight the necessity of studying different mutations in the same gene and have significant implications for future genome editing-based disease modeling studies using hiPSCs and therapeutic interventions for patients with different mutations in the same gene.

## Results

### Differential transcriptomic changes in human hippocampal DGC-like neurons with different heterozygous *SETD1A* mutations

To investigate the functional impact of different *SETD1A* mutations and genetic backgrounds, we first selected 4 hiPSC parental wildtype (WT) lines derived from neurotypical subjects, C7WT^37, 38^, C11WT^39,40^, C14WT, and C15WT^37, 38^, for CRISPR/Cas9 genome editing (Extended Data Table1). For each genetic background, we modeled heterozygous predicted loss-of-function *SETD1A* variants (*SETD1A^+/mut^*) with frameshift mutations targeting either exon 4 (Δe4) or exon 16 (Δe16) (Extended Data Table1). Sanger sequencing showed that indels were the same in Δe16, but different in Δe4 for different hiPSC lines, all of which are predicted to generate premature stop codons in Δe4 or Δe16, respectively (Extended Data Fig. 1a, b). We confirmed the expression of pluripotent cell markers and normal karyotypes of clonally-derived hiPSC lines for 7 isogenic pairs (Extended Data Fig. 1c and 2a).

To obtain a relatively homogeneous population of specific neuron subtypes for characterizing our panel of hiPSC lines, we develop a differentiation protocol that generates a highly pure population of DGC-like neurons. We first patterned feeder-free hiPSC colonies to neural ectoderm in suspension culture as embryoid bodies (EBs) with LDN-193189 and SB-431542 to inhibit the Activin, BMP, and TGFβ signaling pathways for 7 days (Fig. 1a). We then patterned EBs to the dorsal-medial forebrain fate with CHIR-99021 to inhibit GSK3 together with human recombinant BMP7 for an additional 8 days to form human hippocampus-like organoids. We dissociated day 16 organoids into individual neural progenitor cells (NPCs) and cultured them in monolayer with CHIR-99021 for further patterning and proliferation until day 23. Quantitative immunocytochemical analysis at day 23 showed that more than 98% cells were positive for each of the dorsal forebrain neural stem cell markers OTX2, LEF1, LHX2, SOX2, PAX6 and there was no difference between control and mutant lines across different genetic backgrounds (Extended Data Fig. 2b-g). We then dissociated monolayer NPCs on day 23 for culturing with CHIR-99021, B27, BDNF and GDNF until day 37, and finally changed to neuronal maturation medium without CHIR-99021 until day 51 to obtain 4-week-old neurons. Quantitative analysis of 4-week-old neurons showed that more than 96% of the cells were positive for neuronal marker MAP2AB and hippocampal marker ZBTB20 (Extended Data Fig. 3a-c) and more than 96% of the cells were positive for DGC marker PROX1 for each line (Fig. 1b, c). To assess the similarity of our DGC-like neurons to *in vivo* counterparts at the whole transcriptome level, we performed RNA-seq of neuronal cultures and compared them to transcriptomic profiles of neuronal subtypes in subregions of the fetal human hippocampus, including DGCs, CA1 neurons, CA2-4 neurons, subiculum and entorhinal cortex neurons, and GABAergic neurons^41^. We found that transcriptomes of our hippocampal DGC-like neurons were highly correlated with those of DGCs compared to other neuronal subtypes in the human fetal hippocampus (Extended Data Fig. 3d). Together, these results indicate that our differentiation protocol is very robust to reliably generate DGC-like neurons from multiple hiPSC lines and, importantly, there is no difference in the differentiation efficacy between isogenic WT and mutant hiPSC lines from different genetic backgrounds.

**Fig. 1.**
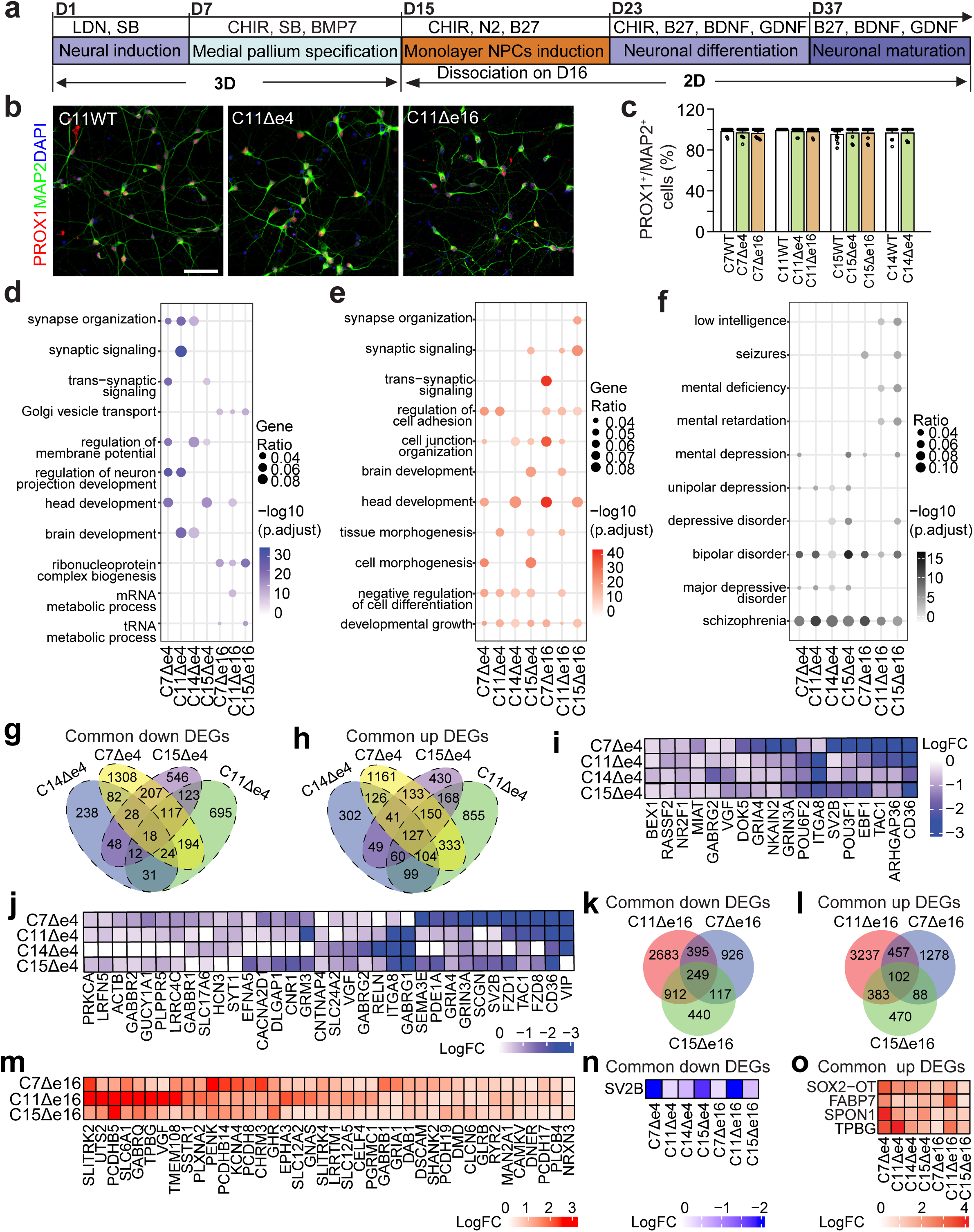
Hippocampal DGC-like neuron differentiation and transcriptomic changes in SETD1A^+/mut^ DGC-like neurons. **a,** Schematic timeline for generating DGC-like neurons from hiPSCs. **b,** Sample confocal images of immunostaining of neuronal marker MAP2 and hippocampal dentate granule cell marker PROX1 in 4-week-old neurons. Scale bar: 50 μm. **c,** Quantification of immunostaining showing the ratio of PROX1 to MAP2 in each line. **d,e,** Gene ontology (GO) analysis of downregulated (**d**) and upregulated (**e**) genes in 4-week-old neurons in each mutant line. **f,** Disease ontology analysis showing disease association in each mutant line. **g, h,** Venn diagrams showing overlap of downregulated and upregulated DEGs in exon 4 mutant lines. **k, l,** Venn diagrams showing overlap of downregulated and upregulated DEGs in exon 16 mutant lines. **i, j,** Heatmap showing common downregulated DEGs (**i**) and downregulated synaptic genes **(j)** in exon 4 mutant lines. **m,** Heatmap showing upregulated synaptic genes in exon 16 mutant lines. **n, o,** Heatmap showing common downregulated and upregulated DEGs among seven mutant lines.

We next characterized *SETD1A* expression in 4-week-old DGC-like neurons derived from the 7 pairs of isogenic *SETD1A^+/mut^*(either Δe4 or Δe16) hiPSC lines on 4 different genetic backgrounds. Comparison of SETD1A protein expression in WT neurons derived from hiPSC lines of 4 genetic backgrounds showed similar levels of expression (Extended Data Fig. 3e). On the other hand, we found a 38-59% reduction of *SETD1A* mRNA by qPCR and a 38-62% reduction of SETD1A protein by western blotting in mutant neurons (Extended Data Fig. 3f-h), which is consistent with the expected dosage effect of heterozygous loss-of-function mutations.

To gain molecular insight into how different *SETD1A* mutations may impact human neurons, we performed bulk RNA-seq analysis of 4-week-old neurons derived from these 7 isogenic pairs cultured in parallel (Extended Data Fig. 4). Comparison of each isogenic pair identified a large set of DEGs (Extended Data Table 2). For mutations targeting exon 4 of *SETD1A* in 4 isogenic pairs, including C7Δe4, C11Δe4, C14Δe4, and C15Δe4, Gene Ontology (GO) analysis of downregulated genes showed enrichment for shared terms related to synapse organization, trans-synaptic signaling, and regulation of membrane potential in at least 2 out 4 pairs (Fig. 1d and Extended Data Fig. 5). For mutations targeting exon 16 of *SETD1A* in 3 isogenic pairs, including C7Δe16, C11Δe16, and C15Δe16, GO analysis of upregulated genes, but not downregulated genes, showed enrichment for terms related to synapse organization, synaptic signaling, and trans-synaptic signaling in some pairs (Fig. 1e and Extended Data Fig. 5). When we combined both up and downregulated DEGs in each pair to perform disease ontology analysis, each mutant line was highly associated with SCZ (Fig. 1f and Extended Data Fig. 5), suggesting a significant role of the *SETD1A* pathway in SCZ pathogenesis in general. We next performed pair by pair comparisons at the individual gene level. We found substantial differences in the upregulated and downregulated DEGs for the same mutation in different genetic backgrounds (Fig. 1g, h, k, l), indicating an important role of the genetic background on the impact of specific mutations on gene expression. In addition, we found 18 common downregulated DEGs (Fig. 1g, i) and 127 common upregulated DEGs (Fig. 1h) among 4 pairs of Δe4 neurons, and 249 common downregulated DEGs (Fig. 1k) and 102 common upregulated DEGs (Fig. 1l) among 3 pairs of Δe16 neurons, suggesting convergence of the impact of specific mutations on gene expression across different genetic backgrounds. In particular, there are a number of common downregulated synaptic DEGs among Δe4 neurons (Fig. 1j) and common upregulated synaptic DEGs among Δe16 neurons (Fig. 1m). There was only 1 common down DEG (SV2B) and 4 common upregulated DEGs (*SOX2-OT, FABP7, SPON1 and TPBG*) among all 7 pairs (Fig. 1n, o). Together, these systematic transcriptomic analyses revealed that the genetic background plays a role in the impact of *SETD1A* mutations at the transcriptome level and similar mutations in a given exon can lead to substantial shared changes across different genetic backgrounds that regulate the same cellular process, such as synapses. More importantly, different mutations of *SETD1A*, all of which were assumed to be loss-of-function mutations, can lead to opposite directional changes in the expression of genes related to the same cellular process across genetic backgrounds.

### Divergent synapse formation and convergent neuronal network changes in different SETD1A^+/mut^ neurons

Given that transcriptomic analysis pointed to changes in the expression of genes related to synapses across all mutant lines, we next examined synapse formation with co-immunostaining of presynaptic marker Synaptotagmin-1 (SYT1) and postsynaptic marker PSD95 for all 7 isogenic pairs of 4-week-old DGC-like neurons. Quantification using the SYN1/PSD95 pair as a synapse marker showed reduced synaptic density in Δe4 neurons, but elevated synaptic density in Δe16 neurons across all 4 genetic backgrounds (Fig. 2a,b), which was consistent with RNA-seq data in which downregulated DEGs were associated with synaptic signaling in Δe4 neurons and upregulated DEGs were associated with synaptic signaling in Δe16 neurons.

**Fig. 2.**
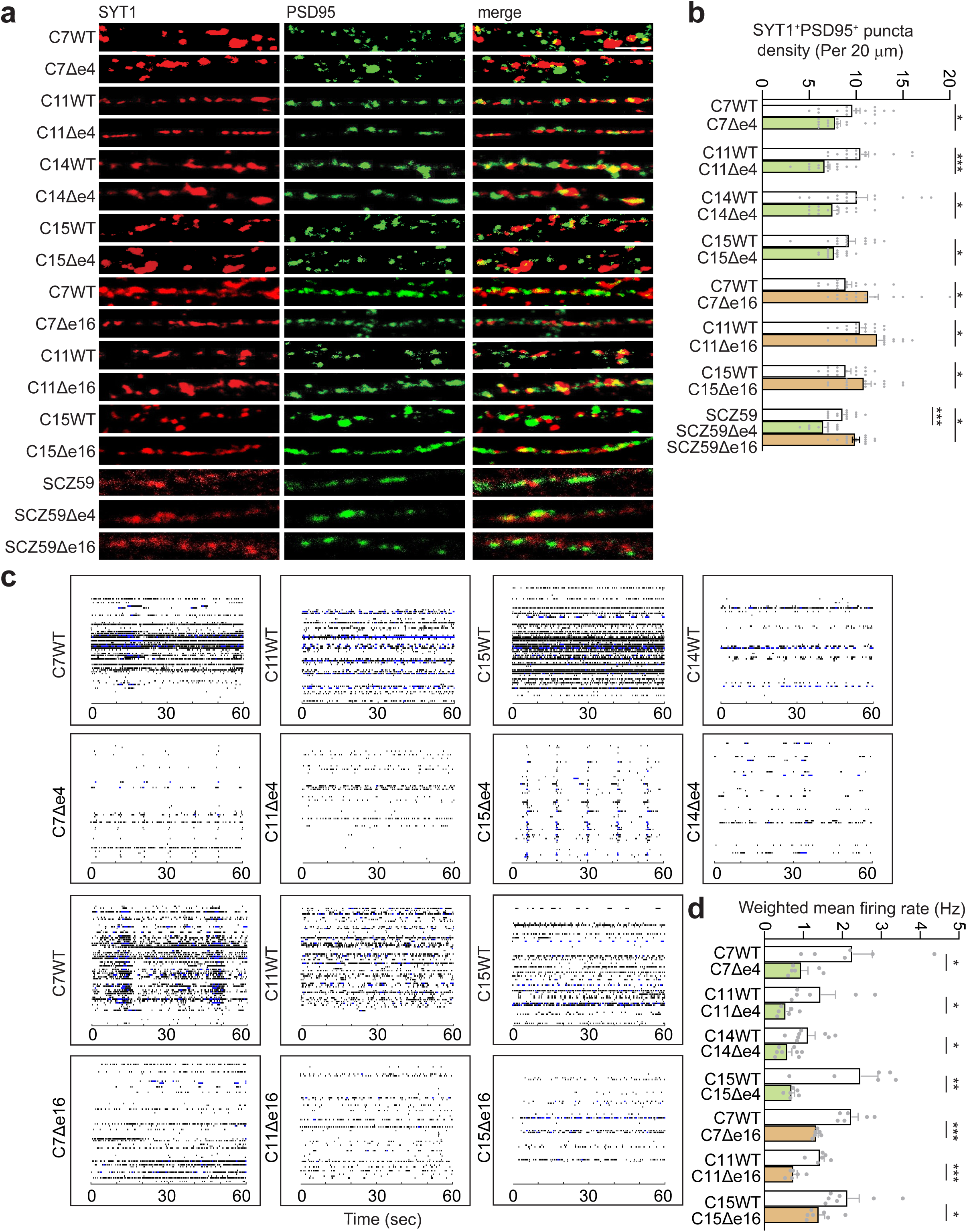
*SETD1A* is essential for synaptic function in hippocampal DGC-like neurons. **a,** Sample confocal images of immunostaining of SYT1 and PSD95 in 9 pairs of 4-week-old DGC-like neurons. Scale bar: 2 μm. **b,** Quantification of co-immunostaining of SYT1^+^ and PSD95^+^ pairs for each pair of neurons. Values represent mean ± s.e.m. (n = 15 neurons derived from 3 independent batches of NPCs in each line, *p < 0.05, **p < 0.01, ***p < 0.001, two-tailed Student’s t-test). **c,** MEA raster plots showing mean firing rate of 7 pairs of 4-week-old DGC-like neurons. **d,** Quantification of weighted mean firing rate for each pair of neurons. Values represent mean ± s.e.m. (n ≥ 6 replicates from 3 independent batches of NPC-derived neurons in each line, *p < 0.05, **p < 0.01, ***p < 0.001, two-tailed Student’s t-test).

To investigate whether a disease-relevant genetic background affects mutation-dependent cellular phenotypes, we engineered exon 4 (SCZ59Δe4) and 16 (SCZ59Δe16) mutations in a hiPSC line derived from one idiopathic SCZ patient^42^ (SCZ59) for a proof-of-principle study and differentiated them into DGC-like neurons with high efficacy similar as for other hiPSC lines in the study (Extended Data Fig. 6). Immunostaining showed that the synaptic density was decreased in SCZ59Δe4 neurons but increased in SCZ59Δe16 neurons at similar levels as in other isogenic pairs (Fig. 2a, b), suggesting that the donor genetic background may not play a major role in affecting how different *SETD1A* mutations affect synapses, perhaps because the highly penetrant mutations have a strong effect.

We further examined the functional synaptic network properties using MEA recordings. For Δe4 neurons, consistent with the decreased number of morphological synapses, spontaneous neuronal firing was significantly lower across different genetic backgrounds. For Δe16 neurons, neuronal firing was also significantly reduced (Fig. 2c, d), which was in contrast to the increased number of morphological synapses and suggests neuronal network-level deficits.

### Transcriptomic, morphological and neuronal network changes with patient mutations

The distinct impacts of different engineered mutations of *SETD1A* on human neurons raise an important question about the effect of patient-specific mutations. We used the C11WT parental line to introduce (via CRISPR/cas9 editing) 2 patient mutations identified in a previous whole exome sequencing study^21^. One mutation of a 4 bp insertion at exon 2 (Y42fs, in 1 patient) generates a premature stop codon (C11KIe2), whereas another mutation of the 2 bp deletion at the exon 16 splice acceptor site (c.4582-2delAG>-; in 4 patients) results in a 79 bp retention of intron 15 with a premature stop codon (C11KIe16) (Extended Data Fig. 7a). The engineered mutations were confirmed by Sanger sequencing, and both hiPSC lines showed normal karyotypes and the expression of pluripotent stem cell markers (Extended Data Fig. 7b-d). Quantitative analysis of immunostaining for dorsal forebrain neural stem cell markers OTX2, LEF1, LHX2, SOX2, PAX6, showed no difference between isogenic hiPSC lines (Extended Data Fig. 7e-j). In 4-week-old DGC-like neurons with patient-specific mutations, over 99% of neurons expressed PROX1 and ZBTB20 in each line (Extended Data Fig. 8a-d). We found a 32-43% reduction of *SETD1A* mRNA and a 34-49% reduction of protein levels, respectively (Extended Data Fig. 8e-g).

We next performed RNA-seq using 4-week-old DGC-like neurons from these 2 pairs of isogenic hiPSC lines. GO analysis of downregulated DEGs in C11KIe2 neurons showed enrichment for terms related to synapse organization, synaptic signaling, and regulation of trans-synaptic signaling (Fig. 3a), resembling those of Δe4 mutant neurons. In contrast, GO analysis of upregulated DEGs in C11KIe16 neurons showed enrichment for terms related to synapse organization and vesicle-mediated transport in synapse (Fig. 3b), similar to Δe16 mutant neurons.

**Fig. 3.**
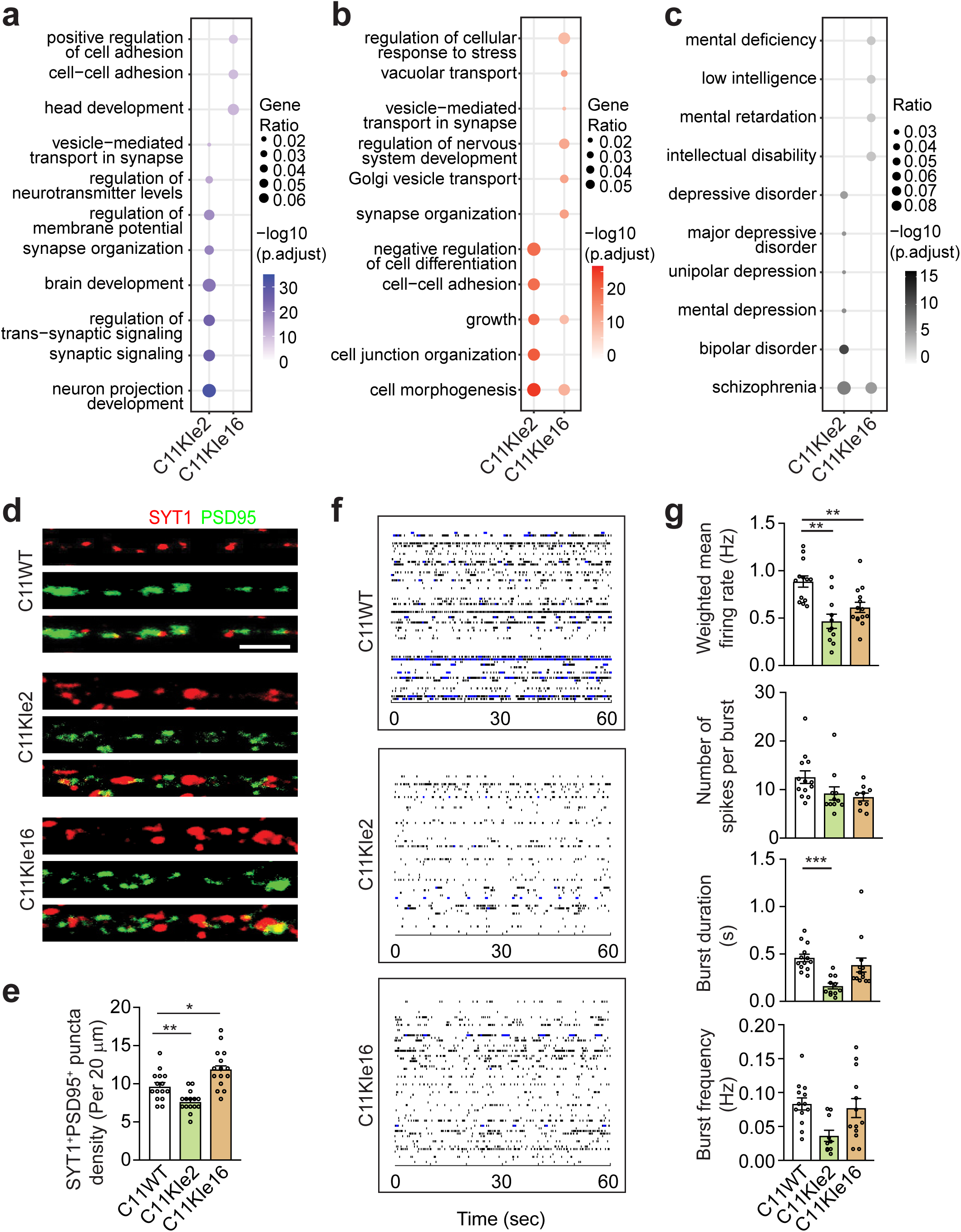
Transcriptomic, morphological, and functional changes in *SETD1A^+/mut^* neurons with patient mutations. **a, b,** GO analysis of downregulated (**a**) and upregulated (**b**) genes in 4-week-old neurons in each patient mutation line. **c,** Disease ontology analysis showing disease association in each mutant line. **d,** Sample confocal images of immunostaining of SYT1 and PSD95 in 4-week-old DGC-like neurons. Scale bar: 2 μm. **e,** Quantification of co-immunostaining of SYT1^+^ and PSD95^+^ pairs for each line. Values represent mean ± s.e.m. (n = 15 neurons derived from 3 independent batches of NPCs, *p < 0.05, **p < 0.01, ***p < 0.001, two-tailed Student’s t-test). **f,** MEA raster plots showing weighted mean firing rate of 4-week-old DGC-like neurons. **g,** Quantification of weighted mean firing rate, number of spikes per burst, burst duration, and burst frequency for each line in 4-week-old DGC-like neurons. Values represent mean ± s.e.m. (n = 11∼14 replicates from 3 independent batches of NPC-derived neurons, *p < 0.05, **p < 0.01, ***p < 0.001, two-tailed Student’s t-test).

Disease ontology analysis of DEGs showed a high association with SCZ for both mutations (Fig. 3c). Furthermore, at both 4 and 8 week time points, C11KIe2 neurons exhibited reduced synaptic density, similar to Δe4 mutant neurons, whereas C11KIe16 neurons, showed elevated synaptic density, similar to Δe16 mutant neurons (Fig. 3d, e and Extended Data Fig. 8h, i). Whole-cell patch-clamp recording of 8-week-old neurons further showed that the mEPSC frequency was decreased in C11KIe2 neurons, but increased in C11KIe16 neurons with no change in mEPSC amplitudes under all conditions, which is in line with our immunostaining results (Extended Data Fig. 8j-l). MEA recordings showed significantly lower neuronal firing in both 4-week-old and 8-week-old neurons from both mutations (Fig. 3f,g and Extended Data Fig. 9), resembling both Δe4 and Δe16 mutant neurons. The discrepancy between the whole-cell patch-clamp assay and the MEA assay suggests that the convergent functional deficits occurred at the level of neuronal network activity.

We had four different mutant lines on the C11WT genetic background. Given that C11Δe4 and C11KIe2 neurons showed similar synaptic and functional phenotypes, we compared RNA-seq data of these two pairs. There were 422 overlapped downregulated DEGs, and GO analysis of these genes showed enrichment for terms related to synaptic signaling and vesicle-mediated transport in synapse (Fig. 4a, b). There were 855 overlapped upregulated DEGs without any synaptic GO terms (Fig. 4c, d). We also compared RNA-seq data of C11Δe16 and C11KIe16 neurons, which showed similar synaptic and functional phenotypes. There were 361 overlapped downregulated DEGs, and GO terms of these genes exhibited enrichment for terms related to modulation of chemical synaptic transmission (Fig. 4e, f). There were 539 overlapped upregulated DEGs, and GO analysis of these genes showed enrichment for terms related to synapse organization and chemical synaptic transmission (Fig. 4g, h). When we extended our analysis to include Δe4 and Δe16 of different genetic backgrounds with pair by pair comparisons, we found 15 common downregulated DEGs among 4 pairs of Δe4 mutant lines and the exon 2 patient mutation line and 34 common upregulated DEGs among 3 pairs of Δe16 mutant lines and exon 16 patient mutation line (Fig. 4i-m).

**Fig. 4.**
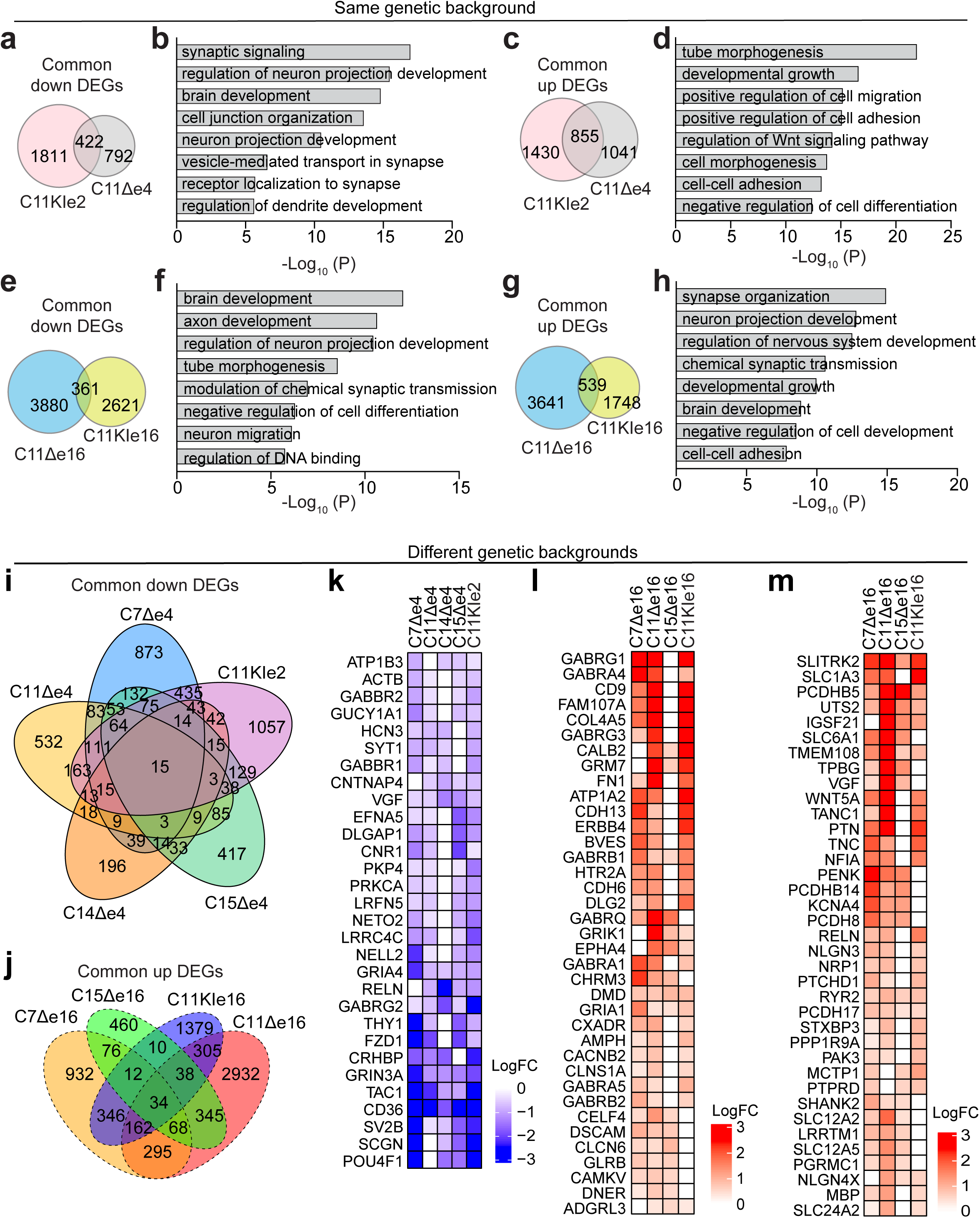
Comparing transcriptomic changes among different hiPSC lines. **a, c,** Venn diagrams showing overlap of downregulated (**a**) and upregulated (**c**) DEGs between C11Δe4 and C11KIe2 neurons. **b, d,** Bar graph showing GO terms of overlap of downregulated (**b**) and upregulated (**d**) DEGs between C11Δe4 and C11KIe2 neurons. **e, g,** Venn diagrams showing overlap of downregulated (**e**) and upregulated (**g)** DEGs between C11Δe16 and C11KIe16 neurons. **f, h,** Bar graph showing GO terms of overlap of downregulated (**f**) and upregulated (**h**) DEGs between C11Δe16 and C11KIe16 neurons. **i,** Venn diagrams showing overlap of common downregulated DEGs among 4 pairs of exon 4 mutant lines plus exon 2 patient mutation line. **j,** Venn diagrams showing overlap of common upregulated DEGs among 3 pairs of exon 16 mutant lines plus exon 16 patient mutation line. **k,** Heatmap showing downregulated synaptic genes in exon 4 and exon 2 mutant lines. **l, m** Heatmap showing upregulated synaptic genes in exon 16 mutant lines.

Together, these results showed substantial differences in gene expression changes induced by different *SETD1A* mutations in distinct genetic backgrounds. Despite this, overall Δe4 mutant neurons appear to resemble the exon 2 patient mutation and Δe16 mutant neurons appear to resemble the exon 16 patient mutation at the molecular, cellular, and functional levels.

### Mechanisms of patient specific mutations

Given the distinct synaptic phenotypes for neurons with early and late exon mutations, we aimed to elucidate the underlying mechanism and we focused on exon 2 and exon 16 patient mutations. We performed a set of “rescue” experiments using lentivirus expressing either full-length WT or exon16 patient-specific mutant (c.4582-2delAG>-) *SETD1A* in different neurons with GFP serving as a reporter of lentiviral infection (Fig. 5a). First, we found that the decreased synaptic density in C11KIe2 neurons was rescued by overexpressing full-length WT *SETD1A* compared to C11WT neurons overexpressing GFP (Fig. 5b, c), suggesting that the exon 2 patient mutation is indeed loss-of-function. Second, we found that the increased synaptic density in C11KIe16 neurons was not rescued by overexpressing full-length WT *SETD1A* compared with C11KIe16 neurons or C11WT neurons overexpressing GFP (Fig. 5b, c), suggesting that the exon 16 patient mutation is not loss-of-function. Finally, we found increased synaptic density in control C11WT neurons overexpressing exon16 patient-specific mutant compared with C11WT neurons overexpressing GFP (Fig. 5b, c), suggesting that the exon 16 patient mutation could be gain-of-function at the phenotypic level.

**Fig. 5.**
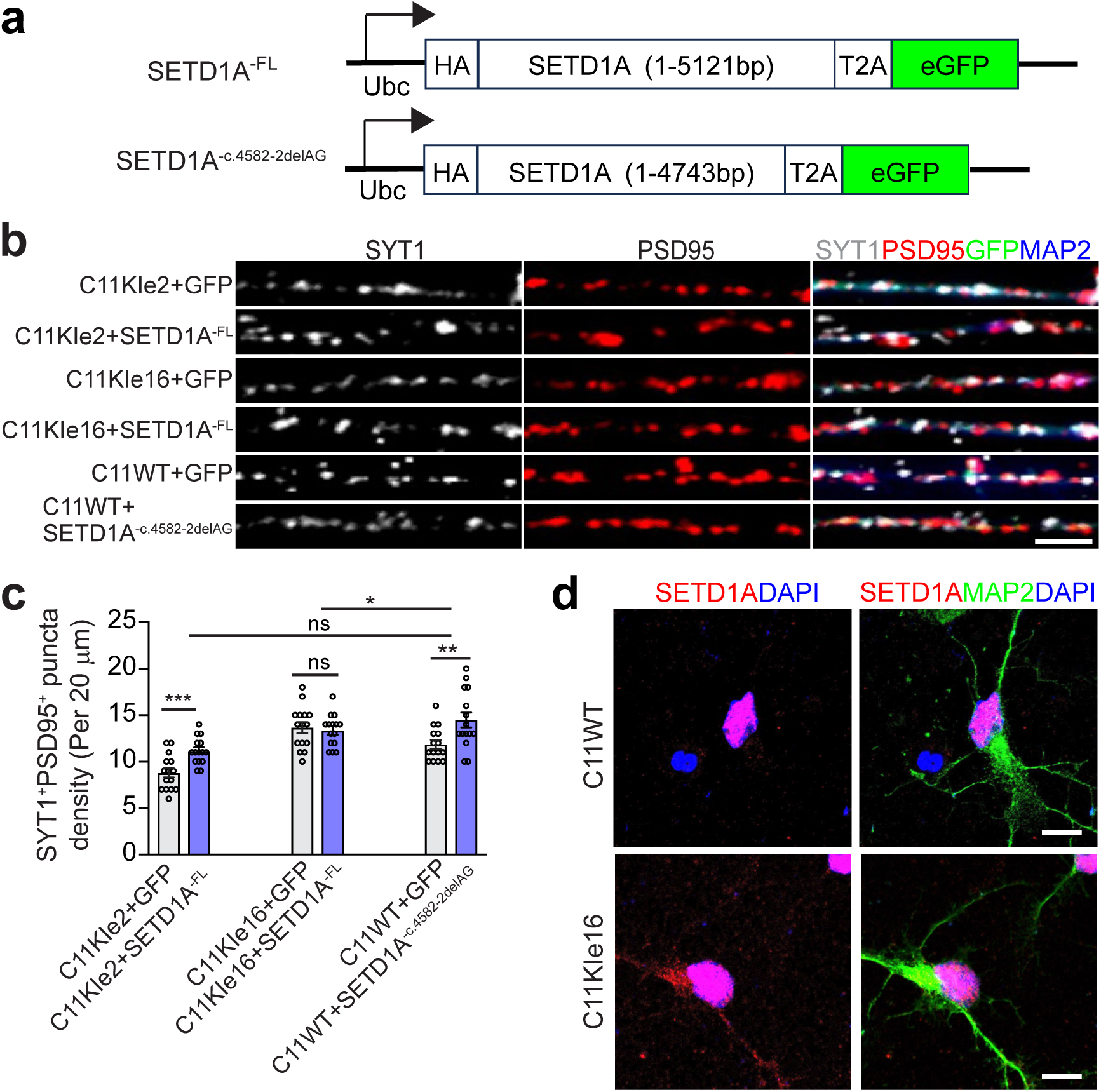
Overexpression of SETD1A constructs in isogenic control and *SETD1A* patient-specific mutation DGC-like neurons and cellular localization of endogenous SETD1A. **a,** Schematic diagram of SETD1A constructs. **b, c,** Representative confocal imaging of synaptic markers, SYT1 and PSD95, and neuronal marker, MAP2, in C11KIe2 and C11KIe16 neurons overexpressing full-length WT SETD1A, and in isogenic control C11WT overexpressing exon 16 patient-specific mutant SETD1A (exon 16 c.4582-2delAG>-) (Scale bar: 2 μm). Neurons overexpressing GFP served as corresponding control (b) and quantification (c). Values represent mean ± s.e.m. (n = 15 infected neurons derived from 3 independent batches of NPCs, *p < 0.05, **p < 0.01, ***p < 0.001, two-tailed Student’s t-test). **d,** Representative confocal images of SETD1A immunostaining using an anti-SETD1A antibody demonstrated nuclear and cytoplasmic localization of endogenous SETD1A protein in C11KIe16 neurons, which are heterozygous for the *SETD1A* exon16 patient-specific mutation (exon 16 c.4582-2delAG>-) (Scale bars: 10 μm).

At the transcriptional level, we found a substantial number of transcripts of the exon 16 patient-specific mutant remained in mutant neurons, while a portion of mutant transcripts was degraded by nonsense-mediated decay (Extended Data Fig. 10a). To examine expression at the protein level, we first expressed HA-tagged full-length SETD1A WT or the exon 16 patient-specific mutant protein in neurons (Extended Data Fig. 10b). Immunostaining of HA revealed that full-length WT SETD1A protein was localized in the nucleus, but exon 16 patient-specific mutant SETD1A protein was localized in the cytoplasm (Extended Data Fig. 10c). To confirm this observation, we performed immunostaining of the endogenous SETD1A protein with an anti-SETD1A antibody in C11WT and C11KIe16 neurons and found that indeed endogenous SETD1A proteins were present exclusively in the nucleus of C11WT neurons, however, endogenous SETD1A proteins were present in both nucleus and cytoplasm of C11KIe16 neurons, which were heterozygous for a *SETD1A* exon 16 patient-specific mutation (Fig. 5d). The differential cellular localization suggests that the exon 16 patient mutation is unlikely to be a dominant-negative mutation.

Together, these results suggested that exon 2 patient-specific mutation represents loss-of-function, whereas exon 16 patient-specific mutation likely represents gain-of-function at the phenotypic level with residual expression of mutation proteins that are mislocalized.

### Different epigenetic impact of *SETD1A* mutations

SETD1A is an H3K4 methyltransferase for the trimethylation of H3K4 (H3K4me3) that generally marks active transcription and is highly enriched at the promoter region^43^. To further understand how different *SETD1A* mutations may affect DGC-like neurons, we performed CUT&RUN for H3K4me3 using 4 pairs of isogenic lines with the same genetic background (C11WT). Indeed, H3K4me3 peaks were enriched in the promoter region in each line (Extended Data Fig. 11a-c). Consistent with reduced levels of SETD1A protein in all these mutant neurons (Extended Data Fig. 3g, h and 8f, g), H3K4me3 read intensity in the TSS region was decreased in each mutant line (Fig. 6a-c). Compared with the paired control lines, each mutant line (C11Δe4, C11KIe2, C11Δe16, and C11KIe16) showed 21797, 25308, 17606, and 22028 decreased H3K4me3 peaks, respectively, mostly in the promoter regions (Extended Data Fig. 11d-g and Extended Data Table 3). To determine whether the reduction in H3K4me3 peaks correlated with transcriptomic changes, we identified overlap of downregulated DEGs from RNA-seq data with genes with decreased peaks in the promoter region from CUT&RUN data. The overlap was 65% and 76% for C11Δe4 and C11KIe2 neurons, respectively, and GO analysis of overlapped genes showed enrichment for terms related to synaptic signaling, regulation of synapse organization, modulation of chemical synaptic transmission, and regulation of trans-synaptic signaling (Fig. 6d, e and h, i). The overlap was lower at 53% and 34% for C11Δe16 and C11KIe16 neurons, respectively, and GO analysis of overlapped genes did not show enrichment for synaptic terms (Fig. 6f, g and j, k). Taken together, these results suggest that while all mutations affect H3K4me3 levels, mutations in exons 2 or 4, which are likely full loss-of-function mutations, lead to a larger impact on gene expression associated with reduction of H3K4me3 compared to exon 16 mutations, especially for synapse-related genes.

**Fig. 6.**
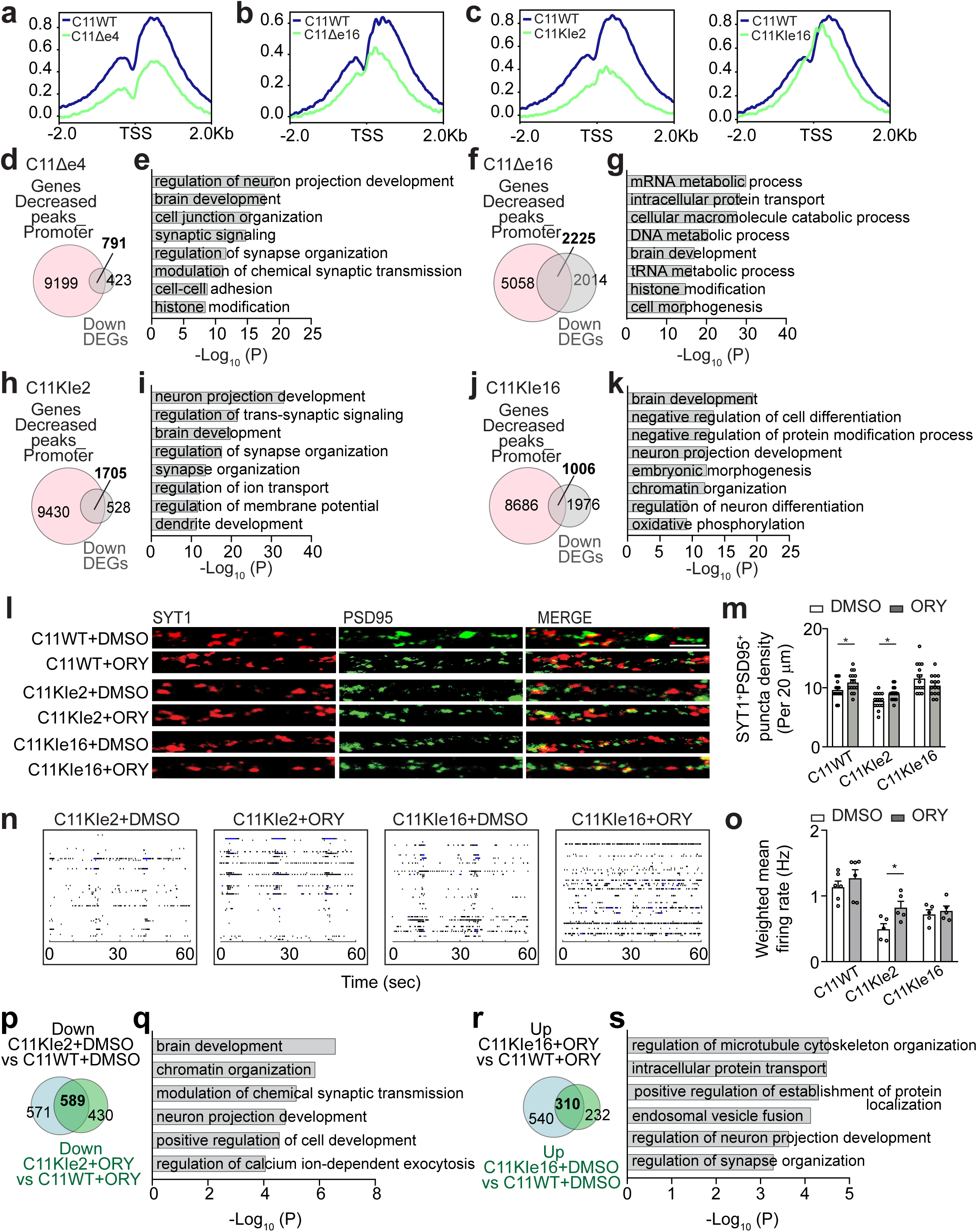
*SETD1A* variants affect global H3K4me3 pattern in human DGC-like neurons, and LSD1 inhibition partially counters *SETD1A* deficiency-induced synaptic deficits only for some mutations. **a, b, c,** Metagene plot of CUT&RUN data showing H3K4me3 reads intensity in the TSS region in one pair of isogenic exon 4 mutant lines (C11WT, C11Δe4), one pair of isogenic exon 16 mutant lines (C11WT, C11Δe16), and 2 pairs of patient mutation lines (C11WT, C11KIe2, C11KIe16). **d, f, h, j**, Venn diagrams showing overlap of downregulated DEGs from RNA-seq data and genes with decreased peaks at promoter regions from CUT&RUN data in each mutant line. e, **g, i, k,** Bar graph showing the GO terms enriched in overlapped genes in d, f, h, j, respectively. **l,** Sample confocal images of immunostaining of SYT1 and PSD95 in 4-week-old DGC-like neurons with DMSO or ORY-1001 treatment. Scale bar: 2 μm. **m,** Quantification of co-immunostaining of SYT1^+^ and PSD95^+^ pair in neurons treated with DMSO or ORY-1001. Values represent mean ± s.e.m. (n = 15 neurons derived from 3 independent batches of NPCs, *p < 0.05, **p < 0.01, ***p < 0.001, two-tailed Student’s t-test). **n,** MEA raster plots showing weighted mean firing rate of 4-week-old DGC-like neurons with DMSO or ORY-1001 treatment. **o,** Quantification of weighted mean firing rate for neurons treated with DMSO or ORY-1001. Values represent mean ± s.e.m. (n = ≥ 5 replicates from 3 independent batches of NPC-derived neurons, *p < 0.05, **p < 0.01, ***p < 0.001, two-tailed Student’s t-test). **p,** Venn diagrams showing downregulated DEGs in C11KIe2 neurons with DMSO or ORY-1001 treatment. **r,** Venn diagrams showing upregulated DEGs in C11KIe16 neurons with DMSO or ORY-1001 treatment. **q, s,** Bar graph showing GO terms enriched in rescued genes in lines C11KIe2 and C11KIe16 after ORY-1001 treatment.

Lysine-specific demethylase 1 (LSD1) is a major demethylase that counteracts the action of SETD1A. Pharmacological inhibition of LSD1 demethylase activity partially rescues the effects of *SETD1A* deficiency in both mouse models^28^ and hiPSC-derived induced glutamatergic/GABAergic neurons^6^. We treated 2-week-old DGC-like neurons derived from two hiPSC lines with patient-specific mutations (C11KIe2 and C11KIe16) with an LSD1 inhibitor, ORY-1001, for 2 weeks. Co-immunostaining of SYT1 and PSD95 and MEA recordings in 4-week-old neurons showed partial rescue of reduced synaptic density and neuronal network properties in C11KIe2 neurons, but not in C11KIe16 neurons (Fig. 6l-o). We also performed RNA-seq to examine transcriptional changes with ORY-1001 treatment. For C11KIe2 neurons, 571 downregulated DEGs were rescued by ORY-1001 treatment, and GO analysis of rescued genes showed enrichment for terms related to modulation of chemical synaptic transmission (Fig. 6p, q). For C11KIe16 neurons, 232 upregulated DEGs were rescued by ORY-1001, and only a few synaptic genes were rescued (Fig. 6r, s).

Together, these findings corroborate results from the set of rescue experiments (Fig. 5) and reveal distinct molecular mechanisms underlying the impact of different mutations in DGC-like neurons at molecular, cellular, and neuronal network levels.

## Discussion

Using a panel of isogenic hiPSC lines with different exon mutations, including engineered premature stop codon mutations in different exons in 5 genetic backgrounds with one idiopathic SCZ patient, and 2 mutations found in SCZ patients on one genetic background, we systematically explored the impact of different mutations and genetic backgrounds on human neurons at molecular, cellular, and neuronal network levels. Our results showed that, while the genetic background can have a substantial impact on *SETD1A* mutation-induced gene expression changes, similar mutations (e.g., in physical proximity) lead to changes in many shared molecular pathways and, more importantly, similarly impact the density of morphological synapses and neuronal network firing across different genetic backgrounds. On the other hand, mutations in different exons can have very different impacts on gene expression and the density of morphological synapses across different genetic backgrounds. Moreover, although not all premature stop codon mutations model loss-of-function, the engineered premature stop codon and patient mutations in proximity or in the same exon may exhibit similar impacts at molecular, cellular, and functional levels. Mechanistically, some of the premature stop codon mutations are indeed loss-of-function, whereas other premature stop codon mutations could potentially be gain-of-function. Together, these results have significant and broad implications for strategies to model genetic risk for disease using engineered iPSC lines.

Patient-derived hiPSCs provide a renewable resource for generating disease-relevant cell types and enable studying the impact of rare genetic variants across different genetic backgrounds. Generating hiPSCs derived from patients with rare genetic variants can be extremely challenging due to limited access to patient donors, but variants can be introduced by CRISPR/Cas9-mediated genome editing^44, 45^. A considerable number of mutations in patients result in a premature stop codon and are predicted to cause loss-of-function^9, 22^. To model such patient mutations with hiPSCs, CRISPR/Cas9 technology is widely used to generate indels that result in premature stop codons. Using *SETD1A* as an example, we generated a large panel of isogenic hiPSC lines to address several important questions related to this general approach in the field.

The first question is whether introduction of premature stop codon mutations in different positions of a gene lead to similar phenotypes. We found clear evidence that the position of the mutation matters and can lead to contrasting molecular and cellular phenotypes that may or may not mimic patient mutations. Notably, premature stop codon mutations in physical proximity appear to cause similar phenotypes. Our results highlight the importance of engineering exact patient mutations for disease modeling, even for mutations predicted to be loss-of-function.

The second question is whether a premature stop codon mutation always leads to loss-of-function. Recent studies of large-scale genome-editing of hiPSCs have shown that not all premature stop codon mutations lead to substantial decreases in the mRNA levels due to nonsense mediated decay^9^. Our finding of the divergent phenotypes between exon 4 and exon 16 as well as 2 patient-specific mutations suggest that not all the *SCZ-*associated rare *SETD1A* mutations that lead to premature stop codons are truly loss-of-function mutations, even if they all lead to an apparent reduction of SETD1A protein and H3K4 methylation levels (Extended Data Fig. 3f, g, Extended Data Fig. 8f, g, and Fig. 6a-c). Our rescue experiment expressing either full length WT or mutant SETD1A in different neurons provides mechanistic insight into the nature of different patient mutations. Specifically, the *SETD1A* patient-specific exon 2 mutation is a *bona fide* loss-of-function mutation. In contrast, the *SETD1A* patient-specific exon 16 mutation, which was presumably “loss-of-function” in previous studies^21, 46^, leads to subcellular mislocalization of endogenous SETD1A mutant protein in human neurons and likely represents a gain-of-function mutation at the phenotypic level, and not a loss-of-function or dominant-negative mutation. In support of this model, an LSD inhibitor partially rescued phenotypes in neurons with the loss-of-function exon 2 patient mutation, but not the gain-of-function exon 16 patient mutation. Importantly, this model is based on evidence obtained from human DGC-like neurons and remains to be examined in other cell types. Notably, *Setd1a* heterozygous mouse models in the C57BL/6 genetic background have been generated in three different laboratories targeting exon 4^28^, exon 7^29^ and exon 15^30^, respectively, and all heterozygous mutations result in *Setd1a* haploinsufficiency. Similar to our findings from human DGC-like neurons, different *Setd1a*-haploinsufficeincy mouse models exhibited differences in behavioral deficits^28–30^ and electrophysiological properties of prefrontal neurons^28, 29^. To date, three hiPSC studies, those by Su et al.^32^, Sawada et al.^31^ and the present study, have engineered a patient-specific *SETD1A* exon 16 mutation (c.4582-2delAG) in hiPSCs with CRISPR and investigated its effects in hiPSC-derived neurons. The study by Su et al. and the present study used one shared parental iPSC line (C11WT), enabling a more direct comparison of findings regardless of influence of the genetic background. Although both studies identified dysregulation of synapse-related genes in mutant neurons, we observed substantially greater transcriptomic changes (present study: 2287 upregulated and 2982 downregulated DEGs; Su et al.: 991 upregulated and 724 downregulated DEGs). Moreover, Su et al. detected no change in the synaptic density, whereas we observed increased synaptic density in mutant neurons. Functionally, both studies demonstrated increased firing rates in mutant neurons using patch-clamp recordings. These discrepancies in molecular and cellular phenotypes could be attributable to two major differences.

First, the studies used different neuronal differentiation strategies. Su et al. employed NGN2-mediated direct neuronal induction, thereby bypassing the NPC stage^32^, whereas our differentiation paradigm proceeds through a NPC intermediate; Different differentiation strategies result in distinct development trajectories, which may contribute to the different neuronal phenotypes^47^. Second, the studies generated neurons with different cellular identities. NGN2-induced neurons assemble fetal cortical neurons^48^, whereas DGC-like neurons resemble hippocampal DGCs in the fetal human brain (Extended Data Fig. 3d). Interestingly, Sawada et al. differentiated cortical neurons through NPCs and observed decreased proliferation and accelerated neurogenesis. However, whether these NPC abnormalities contribute to the subsequent neuronal phenotypes associated with *SETD1A* mutations in Sawada et al. study remains unknown. We focused on the phenotypes of neurons and did not examine NPCs carrying the patient-specific mutation (c.4582-2delAG). Collectively, these studies suggest that findings from hiPSC-based modeling must be interpreted in the context of the specific cell type, developmental stage, experimental paradigm, and genetic background, even if cells carry the same mutation.

The third question is about the impact of the genetic background on phenotypes of different mutations. Our systematic analysis showed that the impact of genetic background and mutations varies depending on the level of analysis. At the molecular level, genetic backgrounds clearly play an important role, where different highly penetrant disease risk mutations can lead to regulation of genes in similar pathways across different genetic backgrounds; at the cellular level, different highly penetrant disease risk mutations could lead to different phenotypes that are consistent for a given mutation across different genetic backgrounds and in our limited study the genetic background of an idiopathic SCZ patient did not appear to alter the effect of *SETD1A* mutations; at the functional level, different highly penetrant disease risk mutations could lead to similar phenotypes across different genetic backgrounds.

When differentiating hiPSCs into a disease-relevant cell type, the composition of the cell population is a primary source of inter-and intra-experimental variation. To date, hippocampal DGC-like neuron differentiation protocols have been reported by two groups. Yu and colleagues reported protocols for both free-floating EB and NPC monolayer approaches, which lead to 35% and 70% 4-week-old PROX1^+^MAP2^+^ DGC-like neurons, respectively, within 68 days^49^. Sakaguchi and colleagues reported a protocol to generate 35% DGC-like neurons over 27 weeks^50^. Robust protocols that produce more homogeneous cell types in a compressed time frame could reduce experimental variability. We developed a robust and efficient protocol to generate highly pure 4-week-old hippocampal DGC-like neurons in 51 days. We confirmed their identity with PROX1 and ZBTB20 immunostaining (Fig. 1b-c, Extended Data Fig. 3a-c) and transcriptomic comparisons showed that our hippocampal DGC-like neurons derived from different genetic backgrounds resemble fetal DGCs (Extended Data Fig. 3d). Our protocol significantly improved the efficiency of generating DGC-like neurons from multiple hiPSC lines. First, our DGC-like neuron differentiation protocol takes a relatively shorter time to generate PROX1^+^MAP2^+^ neurons. Second, the purity (more than 96% in all tested lines with different genetic backgrounds) is much higher than that of cultures generated by published protocols. Third, our protocol can generate a large number of NPCs, which can be greatly expanded over multiple passages and cryopreserved in liquid nitrogen, which makes it possible to use same batch of NPCs across experiments to increase consistency.

Fourth, our DGC-like neurons exhibit functional synaptic transmission without astrocyte co-culture, providing a simplified system for investigating neuron-intrinsic mechanisms and facilitating downstream analyses such as western blotting and high-throughput sequencing. However, the absence of astrocytic support and inhibitory interneurons may limit neuronal maturation and the emergence of synchronized network activity. It will be interesting in the future to examine the impact of S*ETD1A* mutations on properties of astrocytes and how interactions among different cell types may impact phenotypes.

*SETD1A* is one of the top ten SCZ risk genes recently identified to have rare and highly penetrant mutations^22^ and is involved in epigenetic regulation of gene expression through H3K4 methylation to increase the expression of many target genes. Using our novel DGC-like neuron cultures, we comprehensively examined the impact of different mutations of *SETD1A* in 5 different genetic backgrounds on human DGC-like neurons at molecular, cellular, and functional levels. We found that *SETD1A* premature stop mutations in different exons of *SETD1A* resulted in divergent transcriptomic changes with enrichment of downregulated synapse-related genes in exon 2 and exon 4 mutant lines, and conversely, the enrichment of upregulated synapse-related genes in exon 16 mutant lines. Moreover, SCZ is the top disorder associated with DEGs identified from each mutant line. Consistent with the changes at transcriptional levels, immunostaining results show decreased synaptic density in exon 2 and exon 4 mutations, but elevated synaptic density in exon 16 mutations. Despite mutation site-dependent divergent molecular and cellular phenotypes, all mutant lines exhibited convergent deficits in firing rates at the population level (Fig. 2c, d, 3f-g, and Extended Data Fig. 9). Importantly, pharmacological rescue results also suggest a mutation site-dependent response (Fig. 6l-s), suggesting that mutation sites should be considered in the development of treatments for SCZ patients carrying *SETD1A* mutations. Beyond its canonical role as a H3K4 methyltransferase, emerging evidence suggests non-canonical roles of SETD1A. For example, *SETD1A* has been implicated in the DNA damage response through a non-catalytic mechanism involving regulation of DNA repair-associated genes in leukemia cells^51^. More recently, *SETD1A* was shown to bind DNA damage-prone genomic regions in human iPSC-derived neural progenitor cells and neurons, while *SETD1A* haploinsufficiency resulted in increased DNA damage in neurons^31^. Whether *SETD1A* plays non-canonical roles in other cellular processes remains unknown. Future studies investigating non-canonical roles of *SETD1A* and the effects of distinct mutations in different types of human neurons will provide new insights into etiology of SCZ.

In summary, we developed a novel, highly efficient, and reproducible protocol to generate DGC-like neurons and our comprehensive and systematic analysis of different *SETD1A* mutations in these neurons reveal differential impacts of mutation positions and genetic backgrounds on phenotypes at molecular, cellular, and functional levels. Our findings have significant implications for disease modeling using engineered hiPSCs and for precision medicine tailored to patient-specific mutations.

## Methods

### Human iPSC culture and hippocampal DGC-like neuron differentiation

All studies involving hiPSCs were performed under approved protocols of the University of Pennsylvania, Johns Hopkins University and NIH. The source of each hiPSC line is listed in the Extended Data Table 1. All hiPSC lines were confirmed to have a normal karyotype using RNA-seq-based eSNP-Karyotyping^52, 53^ and were confirmed free of mycoplasma with a detection limit of 10 CFU/ml by targeted PCR (Biological Industries). All hiPSC lines were maintained in feeder-free conditions on plates pre-coated with hESC-qualified Matrigel (Corning) using mTeSR plus medium (Stemcell Technologies) as previously described^54, 55^. The hippocampal DGC-like neuron differentiation protocol is illustrated in Fig. 1a. At 70-80% confluence (Day 0), hiPSCs were detached using ReleSR reagent (Stemcell Technologies) and gently pipetted up and down to dissociate colonies. hiPSC clusters were centrifuged at 300 g for 3 minutes and the supernatant was aspirated and replaced with mTeSR Plus medium supplemented with 10 μM Y-27632.

Approximately 3x10^6^ cells were added to an Aggrewell 800 plate (Stemcell Technologies) and cultured overnight in 2ml mTeSR Plus medium supplemented with 10 μM Y-27632 to generate embryoid bodies (EBs). The next day (Day 1), EBs were gently removed from the Aggrewell plate using a P1000 pipettor with a wide-bore tip, washed three times with DMEM:F12, and transferred to neural induction medium (H1 medium, Extended Data Table 1) containing DMEM:F12 supplemented with 20% KnockOut Serum Replacement, 1X MEM-NEAAs, 1X GlutaMax, 1X Penicillin-Streptomycin, 1X 2-mercaptoethanol, 1 μM LDN-193189, 5 μM SB-431542, and 10 μM Y-27632 in ultra-low attachment 6-well plates with 4 mL of medium per well. On day 7, EBs were transferred to ultra-low attachment 6-well plates with 4 mL medial pallium specification medium (H2 medium) containing DMEM:F12 supplemented with 1X N2 supplement, 1X Penicillin/ Streptomycin, 1X MEM-NEAAs, 1X GlutaMax, 3 μM CHIR-99021, 1 μM SB-431542, and 5 ng/mL human recombinant BMP-7 per well. From this point onward, organoids were cultured on an orbital shaker rotating at 110 rpm. On day 16, organoids are dissociated with Accutase at 37°C for 10min and plated onto Matrigel-coated plates in hippocampal DG NPC induction medium (H3 medium) containing a 1:1 mixture of DMEM/F12 and Neurobasal supplemented with 1X N2 supplement, 1X B27 supplement w/o vitamin A, 1X Penicillin/Streptomycin, 1X MEM-NEAAs, 1X GlutaMax and 3 μM CHIR-99021 for one week. On day 23, for neuronal differentiation, cells were dissociated with Accutase and plated onto PDL (or PEI)/ laminin-coated plates or coverslips in neuronal differentiation medium (H4a medium) containing Neurobasal supplemented with 1X B27 supplement, 1X GlutaMax, 200 ng/ml L-Ascorbic Acid, 20 ng/ml BDNF, 20 ng/ml GDNF and 3 μM CHIR-99021. Half of the medium was replaced every 4 days until day 37. Thereafter, neuronal differentiation medium was replaced by neuronal maturation medium (H4b medium) containing Neurobasal supplemented with 1X B27 supplement, 1X GlutaMax, 200 ng/ml L-Ascorbic Acid, 20 ng/ml BDNF and 20 ng/ml GDNF. Half of the medium was replaced every 4-5 days during continuous culturing until day 51.

### *SETD1A* heterozygous generation in hiPSCs with the CRISPR/Cas9 system

Lines C14WT and C14Δe4 were generated in Dr. Zou’s lab. Lines C7WT, C7Δe16, C15WT, and C15Δe16 were generated in Dr. Duan’s lab following a previously published protocol^56^. Briefly, gRNA for SETD1A editing was cloned into the pSpCas9(BB)-2A-Puro (PX459) V2.0 vector (Addgene, # 62988). For editing, 24 hr before transfection, 60-70% confluent parental hiPSCs were dissociated using Accutase and replated onto a matrigel-coated 60mm dish at 4.5 × 10^5^ per well in mTeSR™ Plus with 5 μM ROCK inhibitor. The next day, DNA was transfected using LipofectamineSTEM following vendor’s instructions in antibiotic-free mTeSR™ Plus with 5 μM ROCK inhibitor. 24 hr post transfection, cells were selected in mTeSR™ Plus with 0.5 μg/ml puromycin. The media was refreshed with 0.25 μg/ml puromycin at 48 hr post-transfection until 72 hr post-transfection. Cells were maintained in regular mTeSR™ Plus media for 7-10 days until colonies were large enough for genotyping. The genotypes at the mutagenesis site were confirmed by Sanger sequencing. For the C11Δe4 line, a gene editing method was used as previously described^57^. Briefly, the guide RNA primers were designed by Benchling website (https://www.benchling.com/) (Extended Data Table1) and cloned into the cloning vector (Addgene, Plasmid # 41824) with Gibson Assembly Cloning Kit (New England Biolabs, E5510S). 2 x 10^6^ hiPSCs were electroporated with pCas9_GFP (8 μg) (Addgene, Plasmid #44719) and gRNAs (6 μg/each) using a P3 Primary Cell 4D-Nucleofector X Kit (Lonza) and Nucleofector 4D Device (Lonza) following the manufacturer’s instructions. Positive clones were further confirmed by DNA Sanger sequencing. For the C11Δe16, C7Δe4, C15Δe4, C11KIe2, and C11KIe16 lines, plasmid expressing Cas9-mSA fusion protein was generated by subcloning mSA sequence from the pRSET-mSA plasmid to the pET-21a_3xNLS_SpCas9 plasmid^58^. sgRNAs were designed using the CRIPSR online tool (http://chopchop.cbu.uib.no/) (Extended Data Table1) and synthesized by in vitro transcription using MEGAscript™ T7 Transcription Kit (Invitrogen AM1334). In vitro transcription templates were generated by overlapping PCR using a forward primer containing the T7 promoter followed by an sgRNA targeting sequence and reverse primer containing the sgRNA scaffold sequence. For the patient mutation, biotin-tagged single-strand DNA donors were ordered from IDT. The Cas9/sgRNA/DNA donor complex was delivered into hiPSCs by electroporation using a Neon electroporation kit (Invitrogen MPK1096) following the manufacturer’s instructions. Genotype was checked by PCR and Sanger sequencing. All edited clones were confirmed by Sange sequencing to be free of off-target editing at five top-ranking predicted off-target sites (Extended Data Table 1).

### RNA extraction and qPCR

Four-week-old neurons were dissolved in 1ml Trizol (Invitrogen) for 15 minutes and 200 μl chloroform was then added. After centrifugation 12000g for 15 minutes at 4°C, about 400 μl supernatant was carefully taken out and mixed with the same volume of ethanol, and the mixture was transferred to a Zymo-Spin^TM^ IC Column in a collection tube from the kit of Zymo RNA Clean & Concentrator kit (Zymo Research). After centrifugation, 400 μl RNA Prep buffer, 700 μl RNA wash buffer and 400 μl RNA wash buffer were sequentially added to the column with each procedure followed by centrifugation. Finally, 15 μl nuclease free water was added to elute the column to collect the RNA sample. RNA concentration and quality were assessed using a Nanodrop 2000. Subsequently, about 1μg RNA was used for reverse transcription (RT) to synthesize cDNA with the SuperScript III First-Strand Synthesis System (Invitrogen). Quantitative RT-PCR was then performed using TaqMan (Applied Biosystems) and the StepOnePlus Real-Time PCR system (Applied Biosystems). Quantitative levels for all genes were normalized to the housekeeping gene *GAPDH* and expressed relative to the relevant control samples. All primer sequences are listed in Extended Data Table 1.

### RNA-seq library preparation, sequencing, and analyses

RNA-seq libraries of control and *SETD1A* mutant neurons were prepared based on the SMART-seq2 method as previously described with minor modifications^54, 59^. Briefly, for RT, 3.2 μl RNA (100 ng/μl) was combined with 0.25 μl RNase inhibitor (NEB) and 1 μl CDS primer (10 mM, 5’-AAGCAGTGG TATCAACGCAGAGTACT30VN-3’) in an 8-well PCR tube strip and incubated at 70°C for 2 minutes and immediately placed on ice. Then, 2 ml of 5x SMARTScribe RT buffer (TaKaRa), 0.5 ml of DTT (100 mM). 0.3 ml of MgCl2 (200 mM), 1 μl of dNTPs (10 mM), 1 μl of TSO primer (10 μM, 5’-AAGCAGTGGTATCAACGCAGAGTACATrGrGrG-3’) 0.25 μl of RNase inhibitor (NEB), and 0.5 μl SMARTScribe reverse transcriptase (TaKaRa) were added to the reaction. RT was performed under the following conditions: 42°C for 90 minutes, 10 cycles of 50°C for 2 minutes and 42°C for 2 minutes, 70°C for 15 minutes and 4°C indefinitely. For cDNA amplification, 2 μl of the RT reaction was combined with 2.5 μl of 10x Advantage 2 buffer (Takara), 2.5 μl of 2.5mM dNTPs (Takara), 0.25 μl of 10uM IS PCR primer (5’-AAGCAGTGGTATCAACGCAGAGT-3’), 17.25 μl nuclease free water, and 0.5 μl Advantage 2 DNA polymerase (Takara). Thermocycling conditions were as follows: 94°C for 3 minutes, 9 cycle of 94°C for 15s, 65°C for 30s, and 68°C for 6 minutes, 72°C for 10 minutes, and 4°C indefinitely. Amplified cDNA was then purified using 0.8x AMPure XP beads (Beckman Coulter), eluted in nuclease-free water and quantified following the instructions of Qubit dsDNA HS assay kit (Thermo Fisher Scientific). For tagmentation of cDNA, 50 pg cDNA in 1 μl nuclease free water, 2.5 μl 2xTD buffer (20mM Tris/PH 8.0, 10mM MgCl2, and 16% PEG 8000) and 0.5 μl adaptor-loaded Tn5 transposase (Lucigen) were mixed and incubated at 55°C for 15 minutes and the reaction was terminated by adding 1.25 μl 0.2% SDS and incubated at room temperature for 10 minutes. Fragments were amplified by adding 16.75 μl nuclease free water, 1 μl of Nextera i7 primer (10mM), 1 μl of Nextera i5 primer (10mM), and 25ul KAPA HiFi hotstart readymix (EMSCO/FISHER). Thermocycling conditions were as follows: 72°C for 5 minutes, 95°C for 1 minutes, 14 cycles of 95°C for 30s, 55°C for 30s, and 72°C for 30s, 72°C for 1 minute, and 4°C indefinitely. PCR products were then purified twice with 0.8x AMPure XP beads and eluted in nuclease-free water. Samples were quantified by qPCR (KAPA) and pooled at equal molar amounts. The average fragment size of the final library fragment was quantified by bioanalyzer (Agilent), and the final concentration of the library was determined by qPCR (KAPA). About 2.7 pmol DNA was loaded on NextSeq High Output kit (75 cycles, Illumina) and subjected to NextSeq 550 sequencing (Illumina). Raw sequencing data from RNA-seq were demultiplexed with bcl2fastq2 v2.17.1.14 (Illumina) with adaptor trimmed using Trimmomatic v0.32 software^60^. Alignments were performed using STAR v2.5.2a ^61^ to GENCODE human reference genome GRCh38. Multimapping and chimeric alignments were discarded, and uniquely mapped reads were quantified at the gene level and summarized to gene counts using STAR-quantMode (GeneCounts).

Further analyses were performed in R (v3.6.0). Differentially expressed genes (DEGs) were identified using edgeR^62^. To identify DEGs induced by *SETD1A* mutations, we performed stringent pair-by-pair comparison within the same batch. In our RNA-seq assay, each sample included 3-4 biological replicates derived from independent differentiations. DEGs were identified by comparing a mutant line with its corresponding isogenic control line among multiple biological replicates. The same batch means that NPCs derived from 3-4 independent hiPSC differentiations were cultured, differentiated into neurons and harvested, and libraries were prepared and sequenced at the same time. Our novel protocol enables us to generate and cryopreserve a large number of NPCs from independent differentiations, and then differentiate them into DGC-like neurons at the same time for RNA-seq. Identified upregulated and downregulated gene lists were used for Gene Ontology (GO) enrichment using Metascape database^63^. The fold changes of target differentially expressed genes were visualized in heatmaps by “ComplexHeatmap” package^64^ in R. Disease ontology analysis was performed using WebGestalt^65^, which uses a hypergeometric method and Benjamini Hochberg FDR correction. Spearman correlations were calculated using gene expression levels from RNA-seq for all replicates. Only genes that had non-zero expression levels in all samples were included.

Principal component analysis (PCA) was performed using DESeq2’s plotPCA() function to visualize sample clustering^66^.

### Comparing transcriptome similarity using a random forest-based analysis

To evaluate the transcriptomic similarity between *in vitro* hippocampal DGC-like neurons derived from hiPSC lines with different genetic backgrounds and subregions of the human hippocampal formation (Extended Data Fig. 3d), a previously described multi-class random forest classifier (using the R package ‘randomForest’ (v4.6.14)^67, 68^) was trained on a published single-cell RNA sequencing dataset of the prenatal human hippocampus^41^. Neuronal clusters in different subregions of the hippocampal formation from the published single-cell RNA-seq dataset^41^, including dentate granule cells, CA1 neurons, CA2-4 neurons, subiculum and entorhinal cortex neurons, and GABAergic neurons, were identified using established markers^67^ and selected for similarity comparison analysis. Clusters were randomly downsampled to 1,000 cells to ensure a fair comparison across cell types using ‘sample’ function in R. The number of cells from each cluster *k* used as the training set (*N_k_*) was determined by *N_k_* _=_ min (100, 66% of |cells*_k_*_|_). The top 500 highly variable genes were identified using the ‘FindVariableFeatures’ function in the R package ‘Seurat’ (v4.4.0) and were used to train the random forest classifier on *N_k_* _f_or 500 decision trees. The remaining cells from each of the five neuron types were used as the validation set to estimate the accuracy, resulting in an out-of-bag classification error rate of 11.46%. The classifier was then used to map transcriptomes of *in vitro* hippocampal DGC-like neurons derived from individual iPSC lines obtained by bulk-level RNA sequencing.

### Immunocytochemistry

Cells were fixed with 4% paraformaldehyde for 10min at room temperature, and then permeabilized and blocked with 0.3% Triton X-100 and 10% donkey serum in PBS for 1 hour. Samples were incubated with primary antibodies (Extended Data Table1) at 4°C overnight, followed by incubation with secondary antibodies for 2 hours at 4°C. Images were captured with identical settings for parallel cultures using Zeiss LSM 800 confocal microscope and analyzed with ImageJ (NIH). The total dendritic length was measured by ImageJ plugin NeurphologhJ. For analysis of synaptic bouton density, the numbers of SYT1/PSD95 pairs were manually counted. The synaptic density was determined by D (D = total SYT1/PSD95 pair per 20 μm dendritic length).

### MEA analysis

Twelve-well multielectrode array (MEA) plates (Axion Biosystems) were used as previously described with minor modifications^57^. Briefly, wells were coated with PEI and laminin and neurons were plated at a density of 150,000 neurons/well. MEA plates with neurons were cultured at 37°C and 5% CO2, and cells were fed every 4 days with neuronal differentiation medium (BrainPhys Neuronal Medium N2-A & SM1 kit (Stemcell Technologies), 1 X B27, 1 μM cAMP, 200 ng/ml L-Ascorbic acid, 20 ng/ml BDNF, and 20 ng/ml GDNF). Extracellular recordings of spontaneous action potentials were performed in culture medium at 37°C using a Maestro MEA system and AxIS software (Axion Biosystems). Data were sampled at the rate of 12.5 kHz with a hardware frequency bandwidth of 200-5000 Hz. Five-minute recordings were used to calculate average spike rate.

Spike and burst metrics were analyzed and plotted by Neural Metric Tool version 2.2.3.

### Electrophysiological recordings

Whole-cell patch-clamp electrophysiology for cultured neurons was performed as described^69^. For miniature postsynaptic current recording, a Cs-based internal solution was used, which consisted of (in mM): 40 CsCl, 3.5 KCl, 10 HEPES, 0.05 EGTA, 90 K-gluconate, 1.8 NaCl, 1.7 MgCl_2,_ 2 ATP-magnesium, 0.4 GTP-sodium, and 10 phosphocreatine. Miniature excitatory postsynaptic currents (mEPSCs) were observed in voltage-clamp mode with a holding potential of -70 mV in the presence of 1 μM tetrodotoxin (TTX). The bath solution contained (in mM): 140 NaCl, 5 KCl, 10 HEPES, 2 CaCl_2,_ 2 MgCl_2,_ 10 Glucose, pH 7.4. All cell culture recordings were conducted in the whole-cell configuration at room temperature. The recording files are analyzed using Clampfit (11.1).

### Western blot analysis

To extract proteins from whole cells, neurons were lysed in RIPA buffer (150 mM NaCl, 1% Triton X-100, 0.5% sodium deoxycholate, 0.1% SDS; 50 mM Tris, pH 8.0) containing Complete Protease Inhibitor Cocktail (Roche). Samples were placed on ice for 30 min and sonicated briefly. The insoluble fraction was removed by centrifugation at 15,000 rpm for 15 min at 4°C. Protein concentrations were determined using the Bradford assay (BioRad). Equal amounts of protein were separated by 4-12% SDS-PAGE (BioRad), transferred to nitrocellulose membranes (BioRad), and subsequently incubated with primary (Extended Data Table 1) and secondary antibodies and visualized with SuperSignal West Dura Chemiluminescent Substrate (Thermo Scientific).

Quantification of band intensities was performed using ImageJ software (Extended Data Fig. 12).

### Plasmid cloning

To express SETD1A in neurons, the full length and frameshift coding region from the exon 16 patient mutation (including intron of exon 15) were subcloned to pFUGW plasmid (Addgene #14883). HA tag was added to the 5’ end and the T2A sequence was added to the 3’ end of SETD1A during PCR.

### CUT&RUN assay and bioinformatic analyses

The H3K4me3 CUT&RUN samples were prepared with CUT&RUN kit (EpiCypher). Briefly, neurons were collected in a 1.5 ml tube containing wash buffer. Neurons were captured with ConA beads and incubated with H3K4me3 antibody overnight at 4°C in antibody incubation buffer. After unbound antibodies were washed away, pAG-MNase was added and incubated for 10 min at RT. After washing, CaCl_2 w_as added and incubated for 2 hours at 4°C, and then stopped by adding stop buffer. The protein-DNA complexes were released by incubating at 37°C for 10 min. The supernatant was transferred to a new Lo-bound tube. DNA was then purified using a DNA cleanup kit. Sequencing libraries were prepared using the NEBNext Ultra II DNA library preparation kit for Illumina (New England Biolabs) according to the manufacturer’s instructions. Briefly, end repair was conducted at 20°C for 30 min, followed by dA-tailing at 65°C for 30 min. After adaptor ligation, the DNA fragments were purified by 0.9X volume of SPRIselect beads (Beckman Coulter) followed by 14 cycles PCR amplification with NEBNext Ultra II Q5 Master Mix (New England Biolabs). The PCR products were cleaned up with 0.9× volume of SPRIselect beads. The CUT&RUN libraries were quantified using qPCR (KAPA) and pooled at equal molar amounts, and quality control was performed with the Bioanalyzer High Sensitivity DNA Analysis (Agilent). The libraries were sequenced on the NextSeq High Output kit (75 cycles, Illumina). For the analysis of the CUT&RUN data, all the reads were trimmed, and then uniquely mapped to the hg19 reference genome using Bowtie2^70^. Low mapping quality reads were filtered by SAMtools^71^. Peak calling was performed using MACS2 with default parameters^72^. Intersected peaks defined in both replicates were considered as high confidence peaks. To identify decreased H3K4me3 peaks in mutant lines, MACS2 was used with the file of mutant line serving as the “control” file and the file of its corresponding isogenic control line that is cultured in the same batch serving as the “treatment”.

The BAM files were converted to Bigwig format using deepTools^73^. The normalized read densities of merged replicates were visualized using the Integrative Genomics Viewer (IGV). Peak annotation was performed with ChIPseeker^74^. For the generation of profile plots, the deepTools “computeMatrix” and “plotProfile” functions were used^73^. GO analysis was performed on the Metascape website^63^.

### Quantification and statistical analysis

All statistical tests and sample sizes are included in the Figure Legends and text. All data are shown as mean ± s.e.m. as stated in the Figure Legends. In all cases, the p values are represented as follows: ***p < 0.001, **p < 0.01, *p < 0.05, and ns, not statistically significant when p > 0.05. In all cases, the stated ‘‘n’’ value is either separate wells of monolayer cultures or different batches of cultures. No statistical methods were used to pre-determine sample sizes. For all quantifications of immunohistology, the samples being compared were processed in parallel and imaged using the same settings and laser power for confocal microscopy. For quantification analysis of protein bands in Western blot, the images were cropped with ImageJ software, and the image type was switched to 8-bit. After subtracting the background with setting rolling ball radius as 50.0 pixels, the white and black colors of the images were inverted. Then the intensity of each protein band was measured with the ‘‘Measure’’ function of ImageJ, and the sum of intensity of each protein after normalization with that of loading control from the same sample was used for quantification.

## Data availability

- The RNA-seq and CUT & RUN data reported in this study are deposited in GEO: GSE284412 and GSE284413.
- Any additional information required to reanalyze the data reported in this paper is available from the lead contact upon request.

## Acknowledgements

We thank members of Ming and Song laboratories for discussion and for comments, and B. Temsamrit, E. LaNoce, A. Angelucci, and G. Alepa for laboratory support. This work was supported by grants from National Institutes of Health (R01MH125528 to G-l.M. and Z.P., R01MH106575 to J.D., R35NS097370 and R35NS137480 to G-l.M. and R35NS116843 to H.S.), Dr. Miriam and Sheldon G. Adelson Medical Research Foundation (to G.-l.M.), and UCR start-up funds for Y.H. and T.Z.

## Author contributions

Y.H. and T.Z. co-led the project and contributed equally to this work. Y.H. and T.Z. conducted the main experiments, including cell culture, CRISPR/Cas9 gene editing, MEA, immunocytochemistry, RNA-seq, and CUT&RUN. Q.Y., H.Z., K.M. and J.Z.Z. conducted the CRISPR/Cas9 gene editing, Y.Z. and X.T. M. conducted transcriptome similarity analysis. S.X. conducted electrophysiological recordings. J.Y.L conducted RNA-seq-based eSNP-Karyotyping analysis. A.S. contributed samples from an idiopathic SCZ patient. K.C. contributed to MEA analysis. Z.P.P., J.D., H.S., and G.M. conceived and guided the project. Y.H., T.Z., H.S., and G.M. wrote the paper with inputs from all authors.

## Extended Data Figure Legends

**Extended Data Fig. 1.**
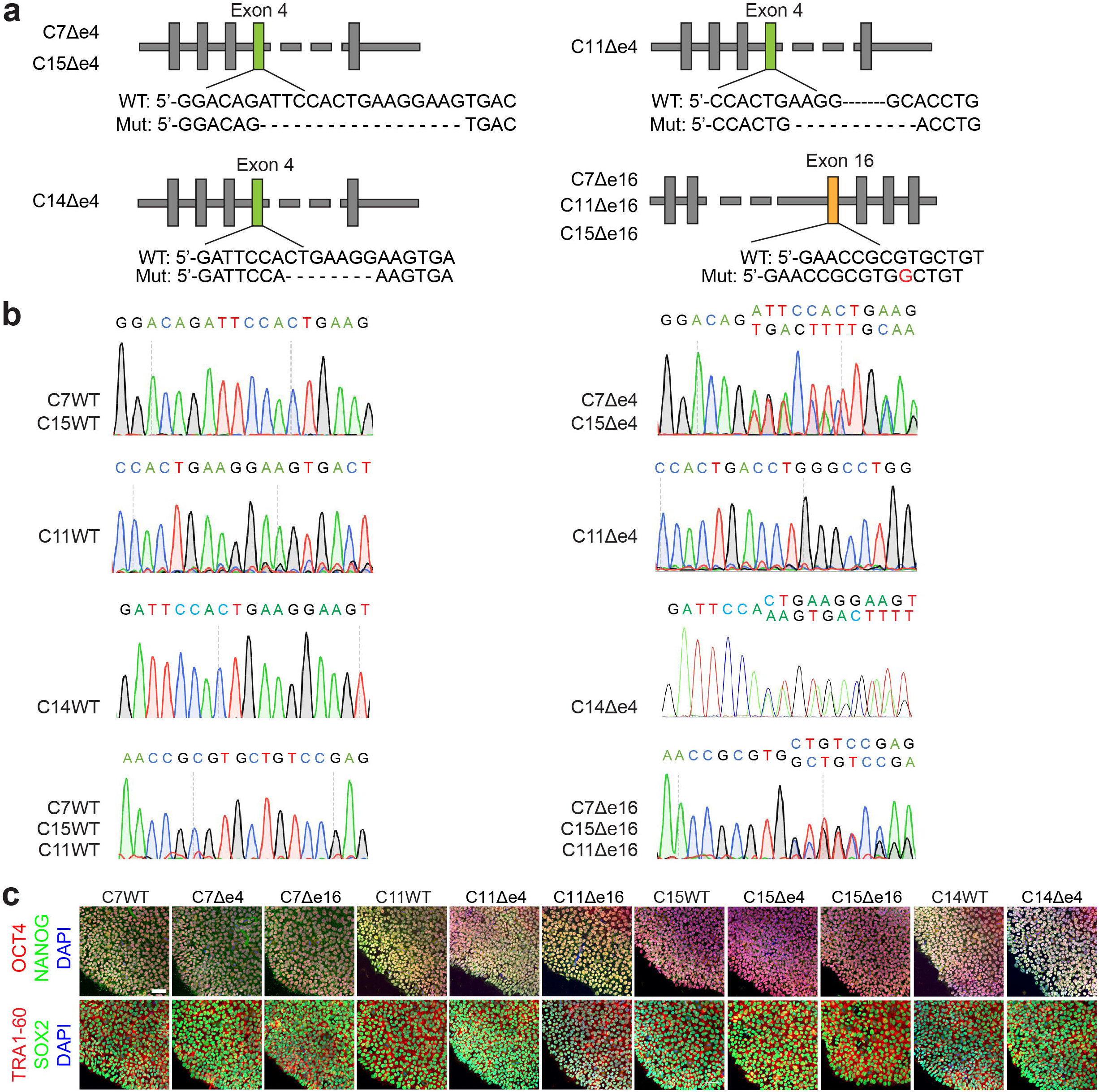
Generation and characterization of *SETD1A^+/mut^* isogenic hiPSC lines using CRISPR/Cas9. **a**, Schematic diagrams showing the mutation sites of each mutant line. **b,** Sanger sequencing trace of the mutation site of each mutant line and corresponding symbol for each line. **c,** Sample confocal images of pluripotency-associated markers for different iPSC lines. Scale bar: 50 μm.

**Extended Data Fig. 2.**
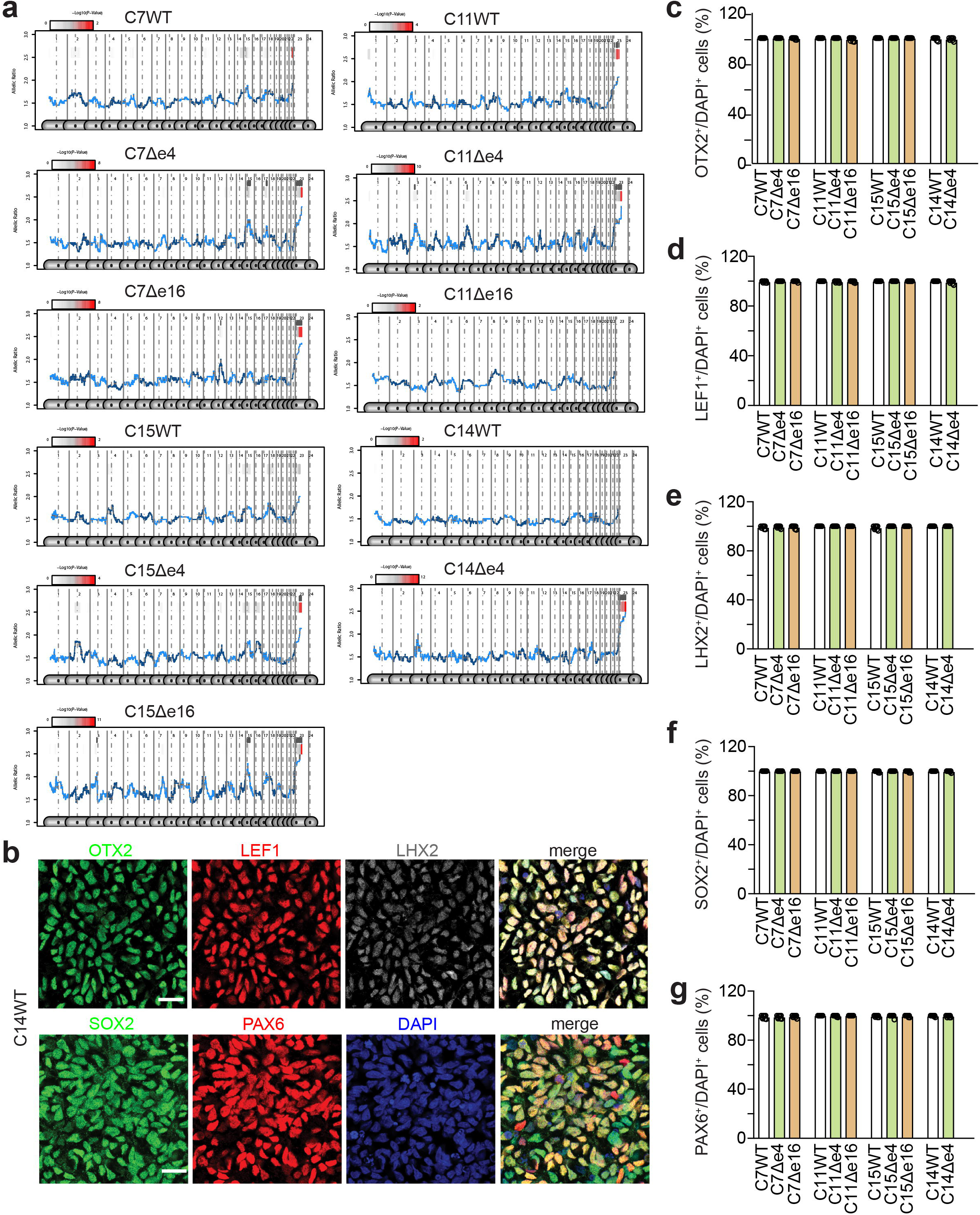
RNA-seq-based eSNP-Karyotyping analysis and characterization of dorsal forebrain neural stem cells. **a,** Each panel shows the moving average of SNP intensity (RNA-seq reads) of the two alleles of heterozygous SNPs. Only the allelic ratio graphs of batch edited lines are shown here; Note that only an autosomal peak interval marked with a red bar above the peak is considered to be significantly “abnormal”, and no hiPSC lines showed such abnormality. **b,** Sample confocal images of immunostaining for dorsal forebrain neural stem cell markers OTX2, LEF1, LHX2, SOX2, and PAX6. Scale bar: 20 μm. **c-g,** Quantification of these markers among different cell lines. Values represent mean ± s.e.m. (n = 4 independent cultures).

**Extended Data Fig. 3.**
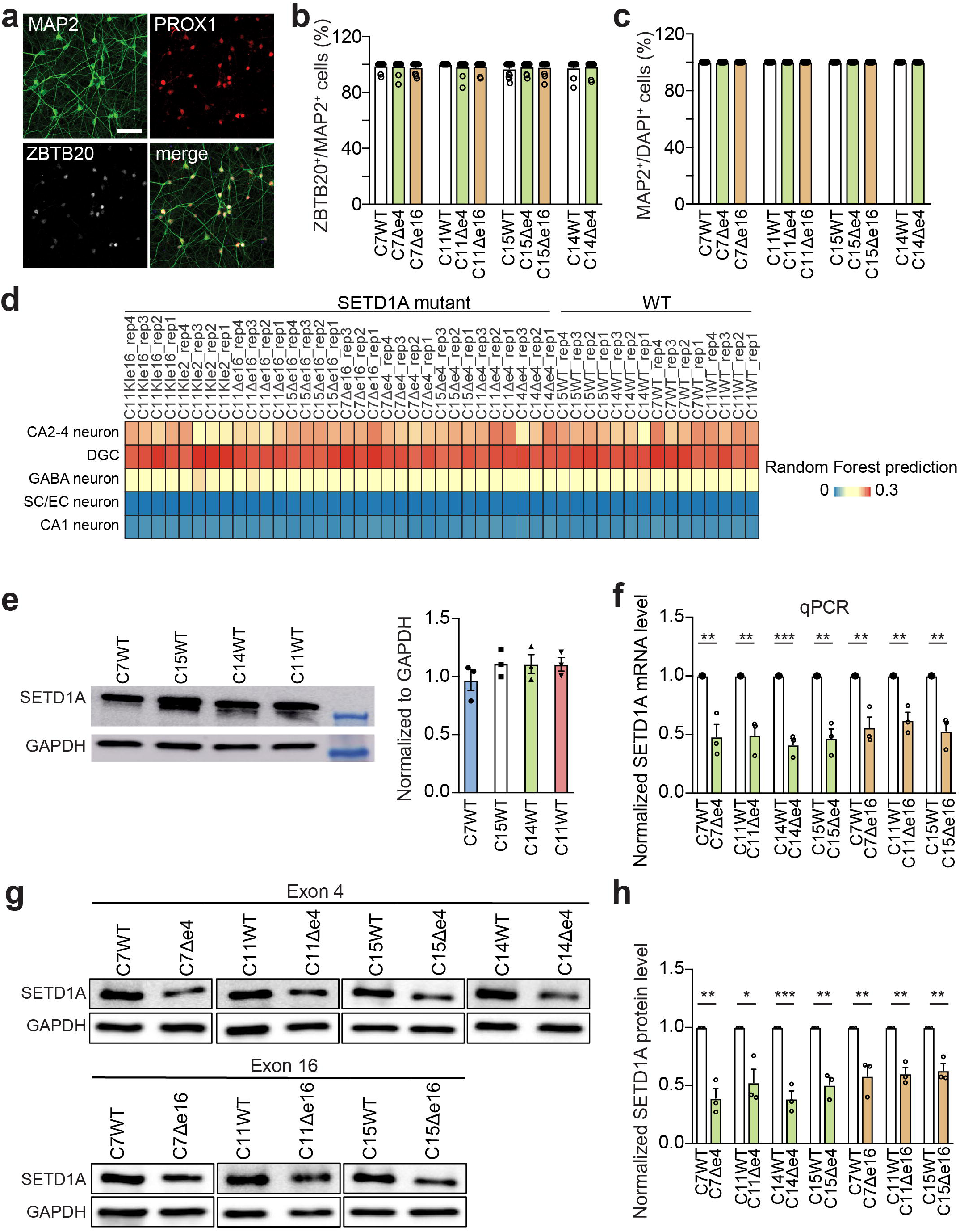
Characterization of human DGC-like neurons derived from *SETD1A^+/mut^*hiPSC lines. **a,** Sample confocal images of immunostaining for neuronal marker MAP2, DG marker PROX1, and hippocampal marker ZBTB20. Scale bar: 50 μm. **b, c,** Quantification of these markers among different cell lines. Values represent mean ± s.e.m. (n = 4 independent cultures). **d,** Transcriptome similarity of *in vitro* DGC-like neurons derived from different genetic backgrounds hiPSC lines and neurons from subregions of the prenatal human hippocampal formation was compared using a random forest-based analysis. **e,** Western blotting and quantification of endogenous SETD1A protein in one-month-old DGC-like neurons derived from four parental lines. Values represent mean ± s.e.m. (n = 3 independent cultures). **g,** Quantification of *SETD1A* mRNA levels from qPCR analysis. Data were normalized to control in each pair. Values represent mean ± s.e.m. (n = 3 cultures, *p < 0.05, **p < 0.01, ***p < 0.001, two-tailed Student’s t-test). **g, h,** Western blotting images and quantification of SETD1A protein levels with anti-N-terminal antibodies. Data were normalized to control in each pair. Values represent mean ± s.e.m. (n = 3 independent cultures, *p < 0.05, **p < 0.01, ***p < 0.001, two-tailed Student’s t-test). **i,** Schematic diagrams showing different mutant *SETD1A* constructs. Western blotting images showing SETD1A protein expression levels of each construct using either HA antibody or N-terminal SETD1A antibody.

**Extended Data Fig. 4.**
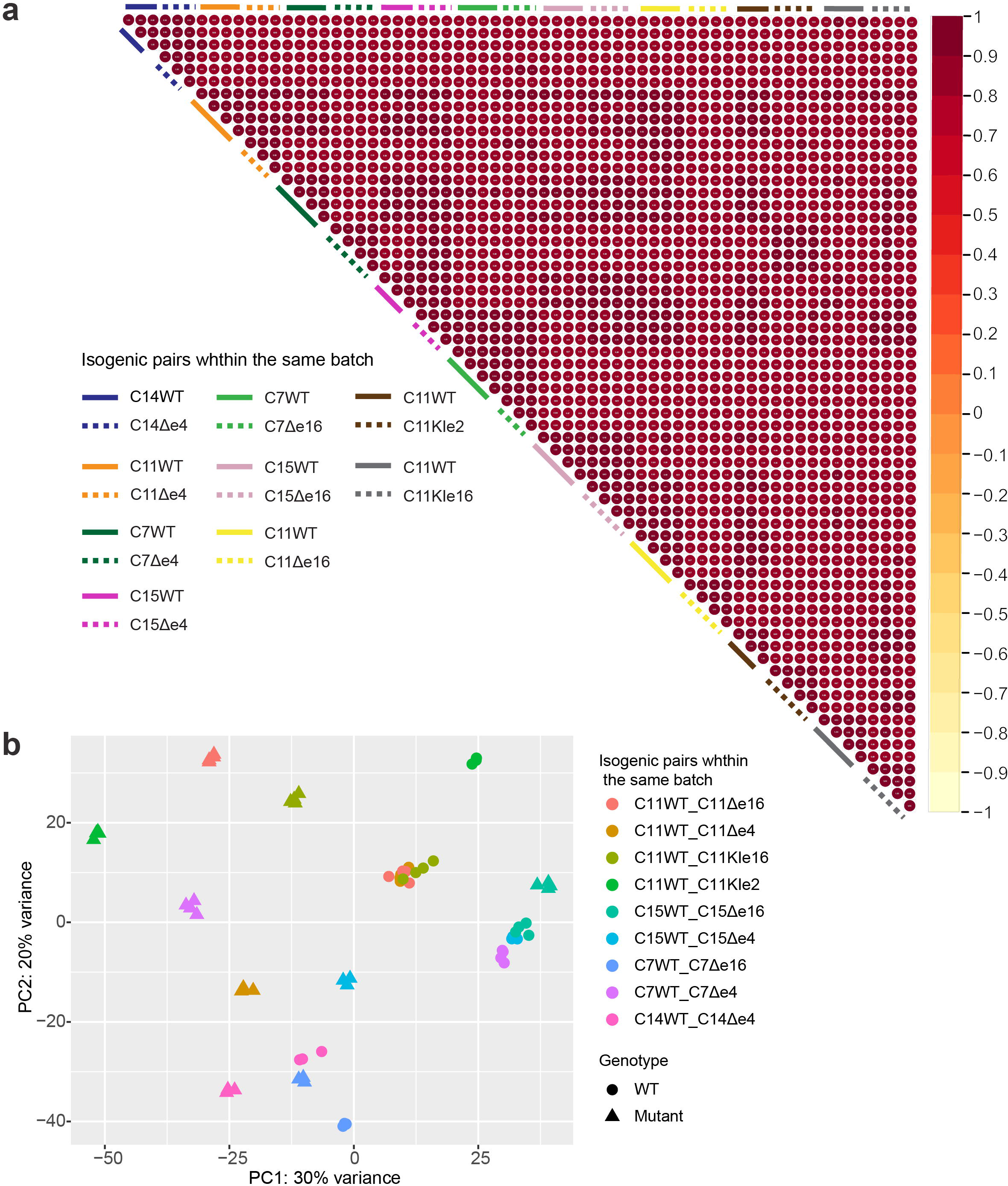
Spearman’s correlation coefficients and the PCA plot across all samples for RNA-seq assay. **a,** Heat map of Spearman’s correlation coefficients across all samples in RNA-seq assay. Each sample includes 3-4 biological replicates underneath the bar. Solid bars represent isogenic controls, and dashed bars represent mutations. Each isogenic pair within the same batch was labelled by one color. **b,** PCA plot showed clusters of all biological replicates for RNA-seq assay.

**Extended Data Fig. 5.**
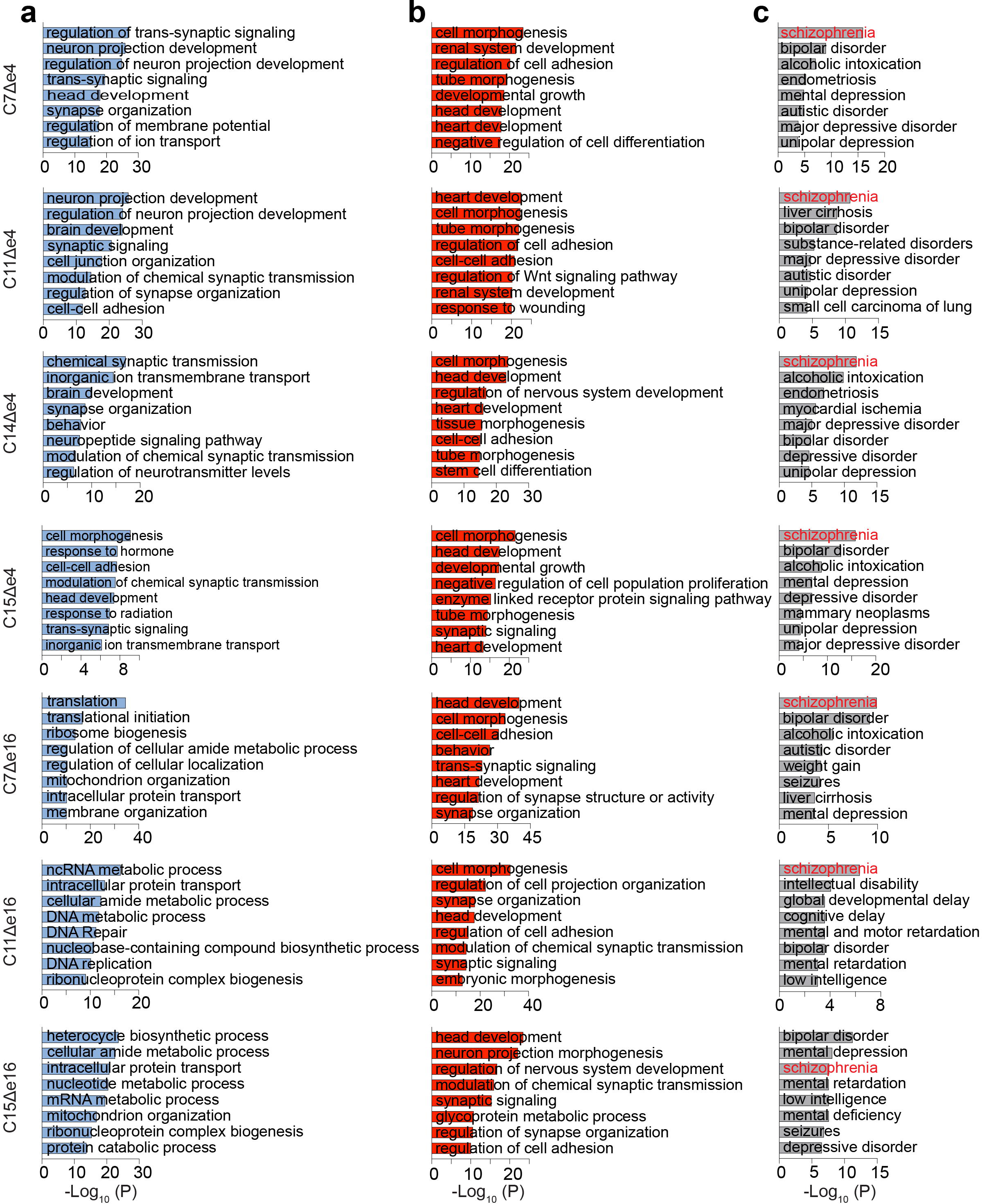
Detailed GO terms and disease ontology analyses. **a, b,** Bar graphs showing detailed GO terms enriched in downregulated (**a**) and upregulated (**b**) DEGs in each mutant line. **c,** Bar graphs showing detailed disease ontology analyses for each mutant line.

**Extended Data Fig. 6.**
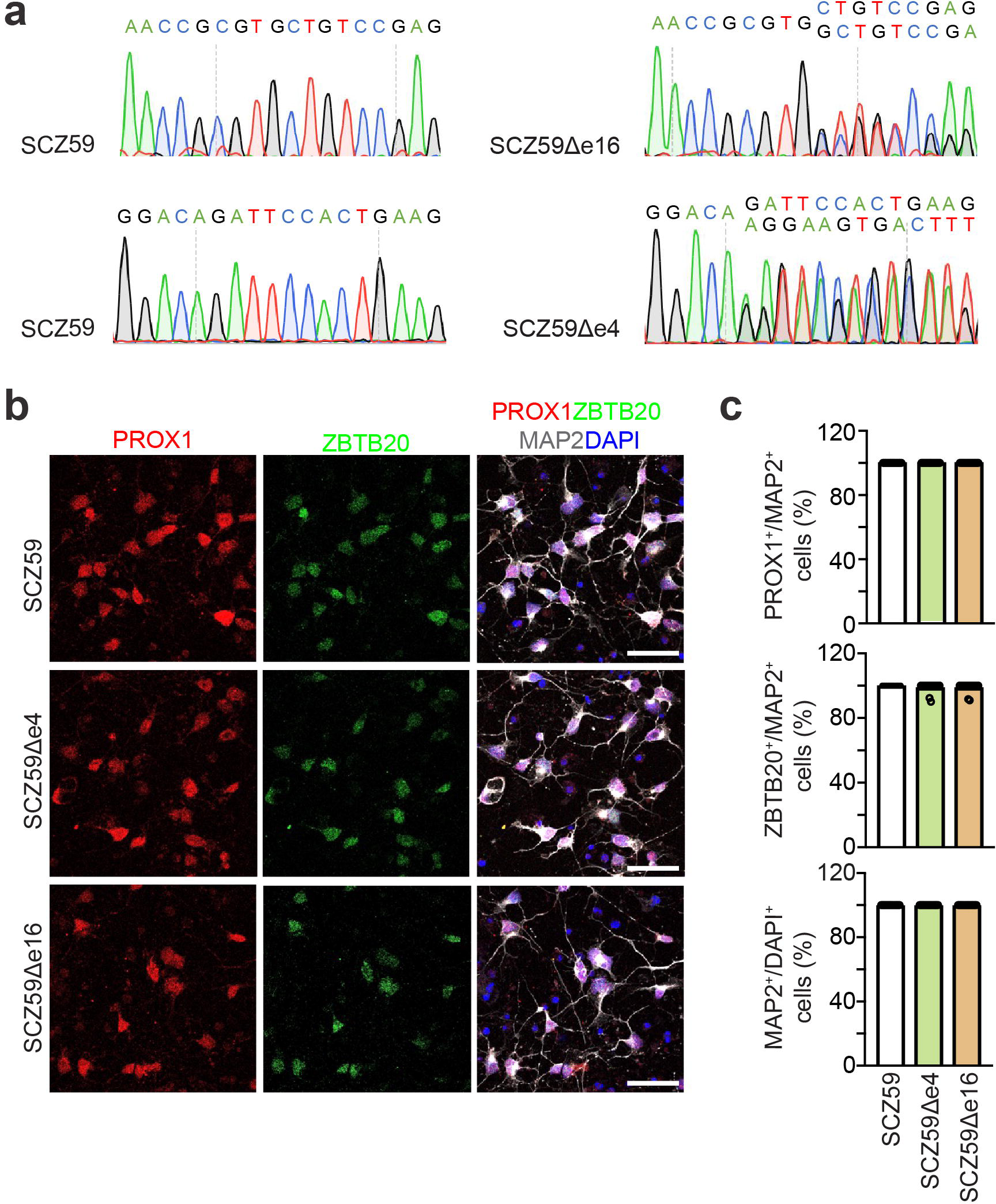
Characterization of DGC-like neurons derived from an hiPSC line from an idiopathic SCZ patient and quantification of the synaptic density. **a** Sanger sequencing trace of mutation site of each mutant line and corresponding symbol for each line. **b, c,** Representative confocal imaging of hippocampal dentate granule neuron markers PROX1 and ZBTB20 (**b**) (Scale bars: 40 μm) and quantification of purity of DGC-like neurons derived from the hiPSC line from a schizophrenia patient, SCZ59 (**c**). Values represent mean ± s.e.m. (n = 15 neurons derived from 3 independent batches of NPCs; *p < 0.05, **p < 0.01, ***p < 0.001, two-tailed Student’s t-test).

**Extended Data Fig. 7.**
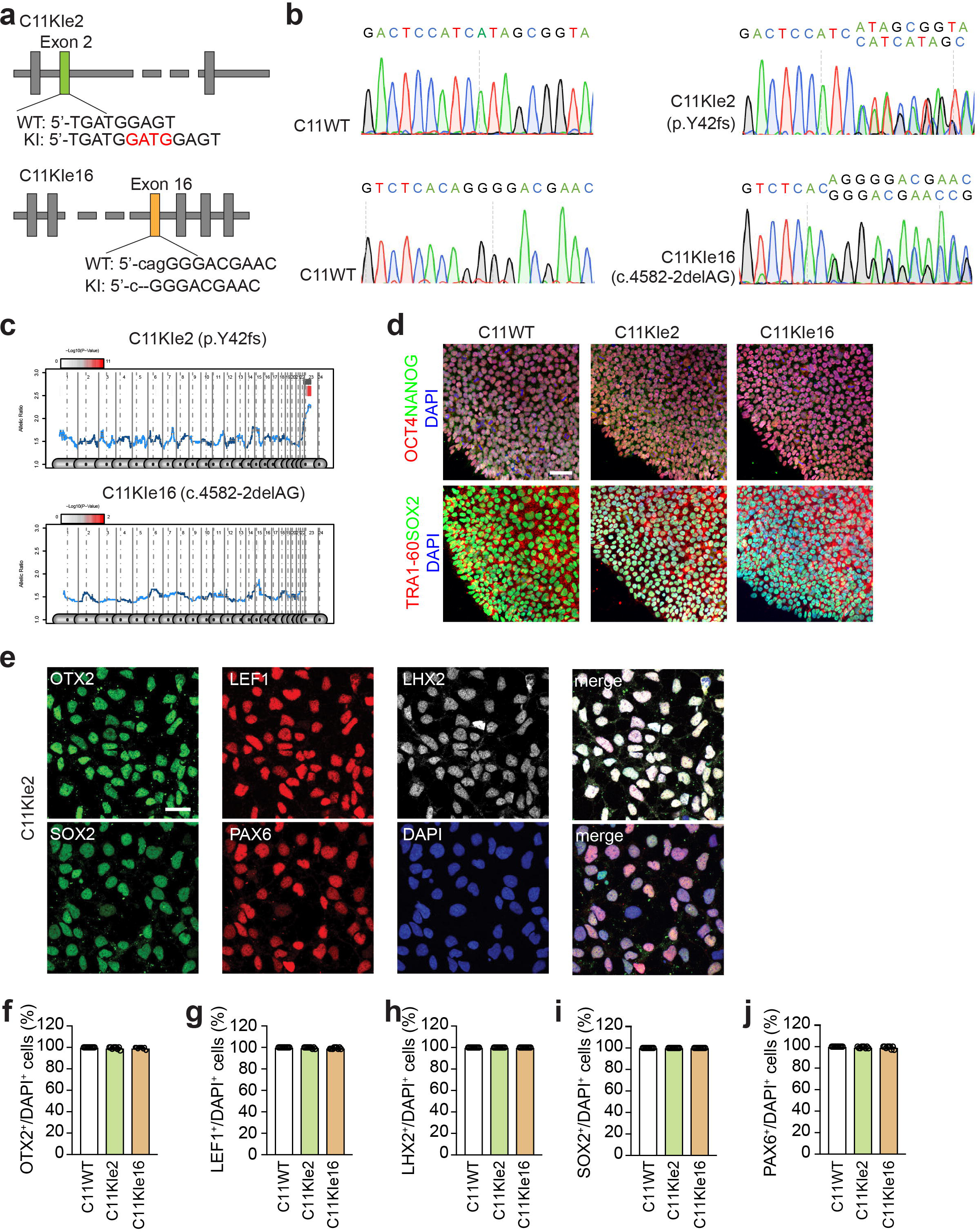
Generation and characterization of hiPSC lines with patient-specific *SETD1A* mutations using CRISPR/Cas9. **a,** Schematic diagrams showing mutation sites of each patient mutation line. **b,** Sanger sequencing trace of the mutation site of each mutant line and corresponding symbol for each line (Sanger sequencing traces for the C11WT and C11KIe2 pair are from 3’-5’). **c,** Each panel shows the moving average of SNP intensity (RNA-seq reads) of the two alleles of heterozygous SNPs. Only the allelic ratio graphs of batch edited lines are shown here; Note that only an autosomal peak interval marked with a red bar above the peak is considered to be significantly “abnormal”, and no h iPSC line showed such abnormality. **d,** Sample confocal images of immunostaining for pluripotency-associated markers for each line. Scale bar: 50 μm. **e,** Sample confocal images of immunostaining for dorsal forebrain neural stem cell markers OTX2, LEF1, LHX2, SOX2, and PAX6. Scale bar: 20 μm. **f-j,** Quantification of these markers among different cell lines. Values represent mean ± s.e.m (n = 4 independent cultures).

**Extended Data Fig. 8.**
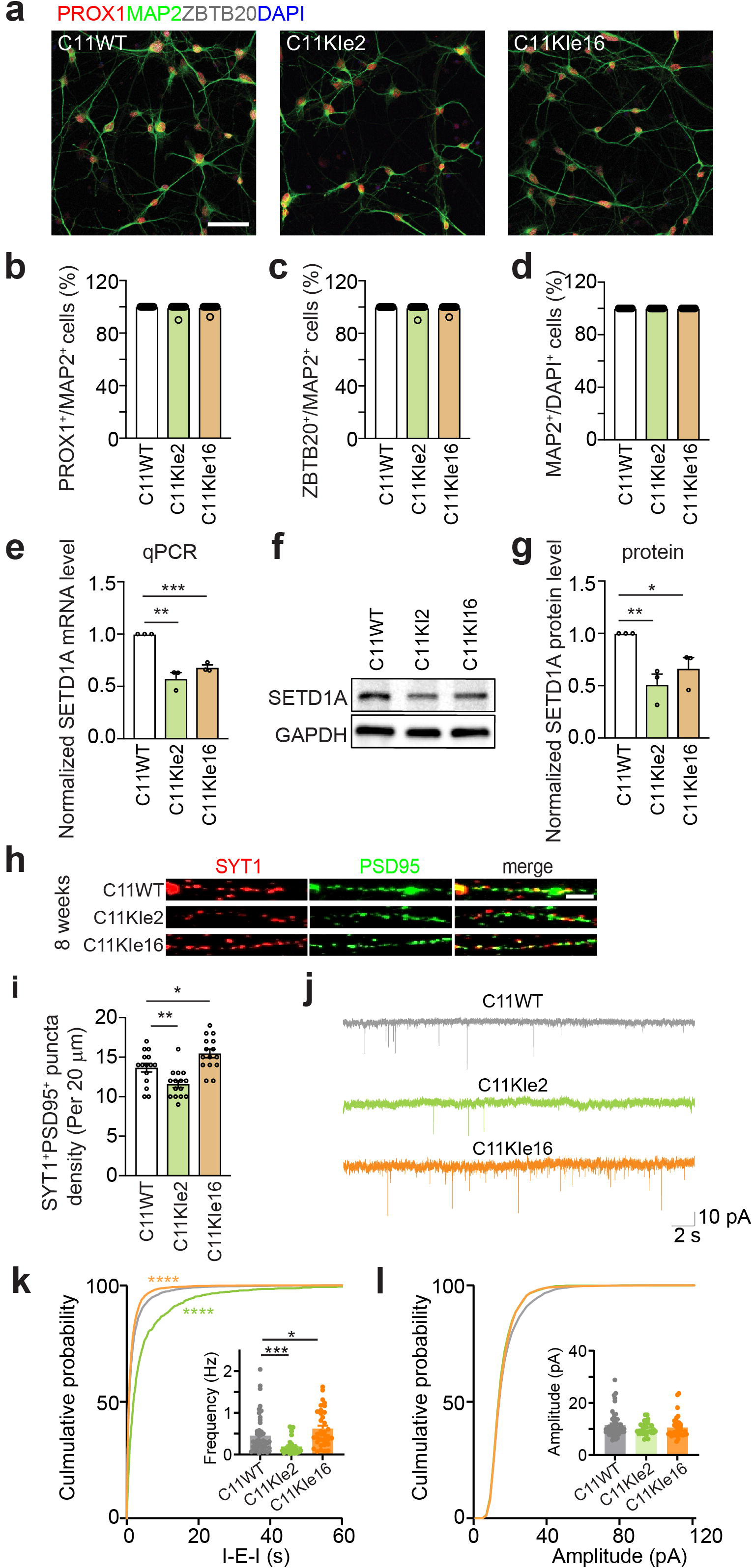
Characterization of human DGC-like neurons with *SETD1A* patient mutations. **a,** Sample confocal images of immunostaining for neuronal marker MAP2, DG marker PROX1, and hippocampal marker ZBTB20. Scale bar: 20 μm. **b-d,** Quantification of these markers among different cell lines. **e,** Quantification of *SETD1A* mRNA levels from qPCR analysis. Data were normalized to control. Values represent mean ± s.e.m. (n = 3 independent cultures, *p < 0.05, **p < 0.01, ***p < 0.001, two-tailed Student’s t-test). **f, g,** Western blotting images and quantification of SETD1A protein levels. Data were normalized to control. Values represent mean ± s.e.m. (n = 3 cultures, *p < 0.05, **p < 0.01, ***p < 0.001, two-tailed Student’s t-test). **h, i**, Sample confocal images of immunostaining of synaptic markers SYT1 and PSD95 in 8-week-old DGC-like neurons (**h**) and quantification (**i**). (Scale bar: 2 μm). Values represent mean ± s.e.m. (n = 15 neurons derived from 3 independent batches of NPCs, *p < 0.05, **p < 0.01, ***p < 0.001, two-tailed Student’s t-test). **j-l**. Patch clamp assay revealed decreased mEPSC frequency in neurons with exon 2 patient-specific mutation, but increased mEPSC frequency in neurons with exon 16 patient-specific mutation (**k**), but no significant difference in the mEPSC amplitude between isogenic control and patient-specific mutations (**l**). Also shown are representative traces of mEPSC recording for C11WT, C11KIe2, and C11KIe16 neurons (**j**). (*p < 0.05, ***p < 0.001; Mann-Whitney test for the frequency and amplitude; Kolmogorov-Smirnov test for the cumulative probability).

**Extended Data Fig. 9.**
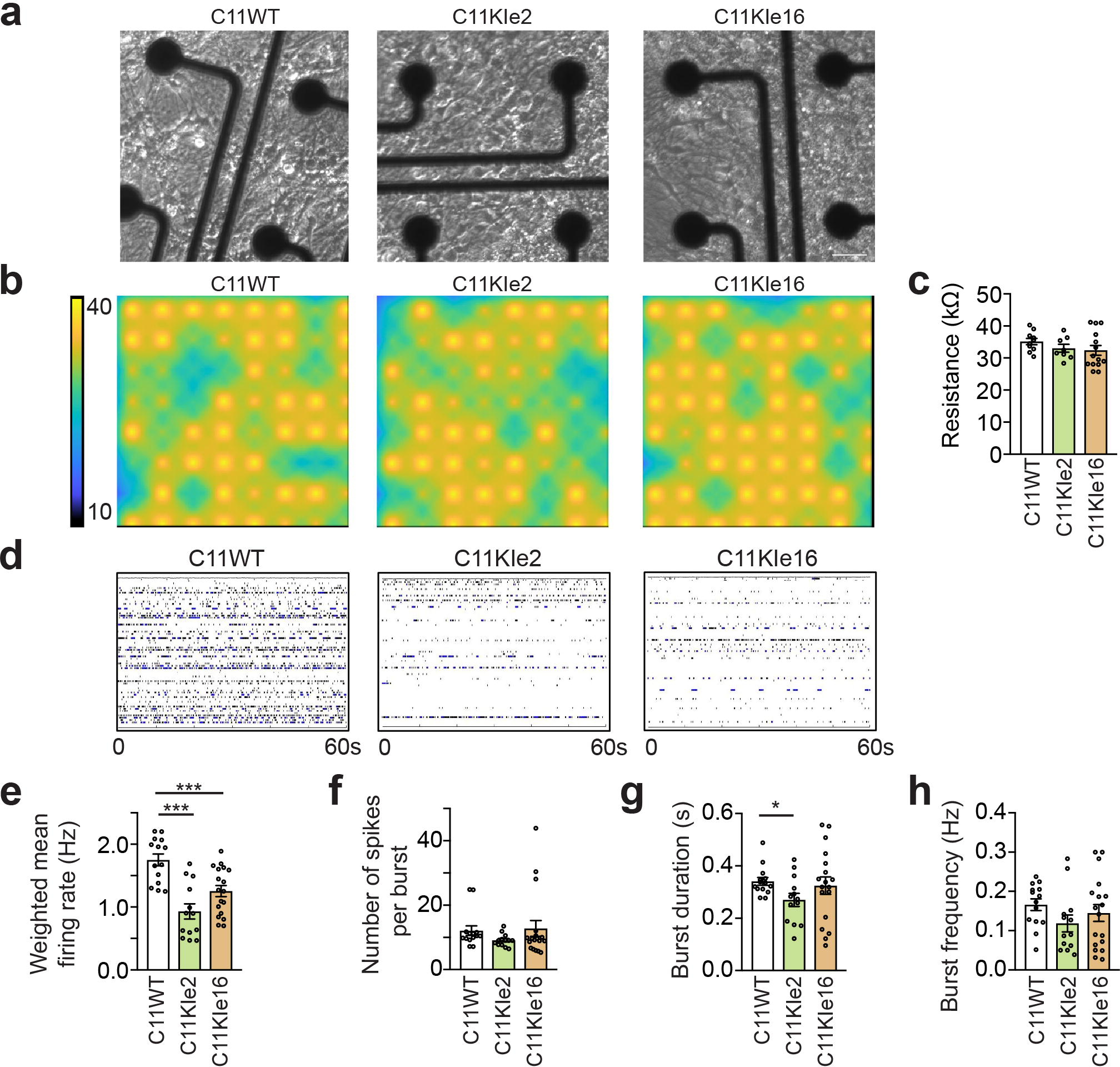
MEA assay of 8-week-old DGC-like neurons carrying different *SETD1A* patient-specific mutations. **a**, Representative images of 8-week-old isogenic control and *SETD1A* exon 2 and exon 16 patient-specific mutation DGC-like neurons in the MEA assay (Scale bar: 50 μm). 1.5 x 10^5^ neurons for each line were plated in each well of 12-well MEA plates. **b**, Representative heatmap for viability of 8-week-old isogenic control and *SETD1A* exon 2 and exon 16 mutation DGC-like neurons on the array. Each dot represents an electrode (64 electrodes per well). **c**, Resistance is a measure of viable cell coverage over the electrode. Higher values indicate more intact cells are attached to the electrode. No significant differences were observed in resistance between isogenic control and mutation lines. **d,** MEA raster plots showing weighted mean firing rates of 8-week-old DGC-like neurons. **e-h,** Quantification of weighted mean firing rates, number of spikes per burst, burst duration, and burst frequency for each line. Values represent mean ± s.e.m. (n = 14-17 replicates from 3 independent batches of NPC-derived neurons, *p < 0.05, **p < 0.01, ***p < 0.001, two-tailed Student’s t-test).

**Extended Data Fig. 10.**
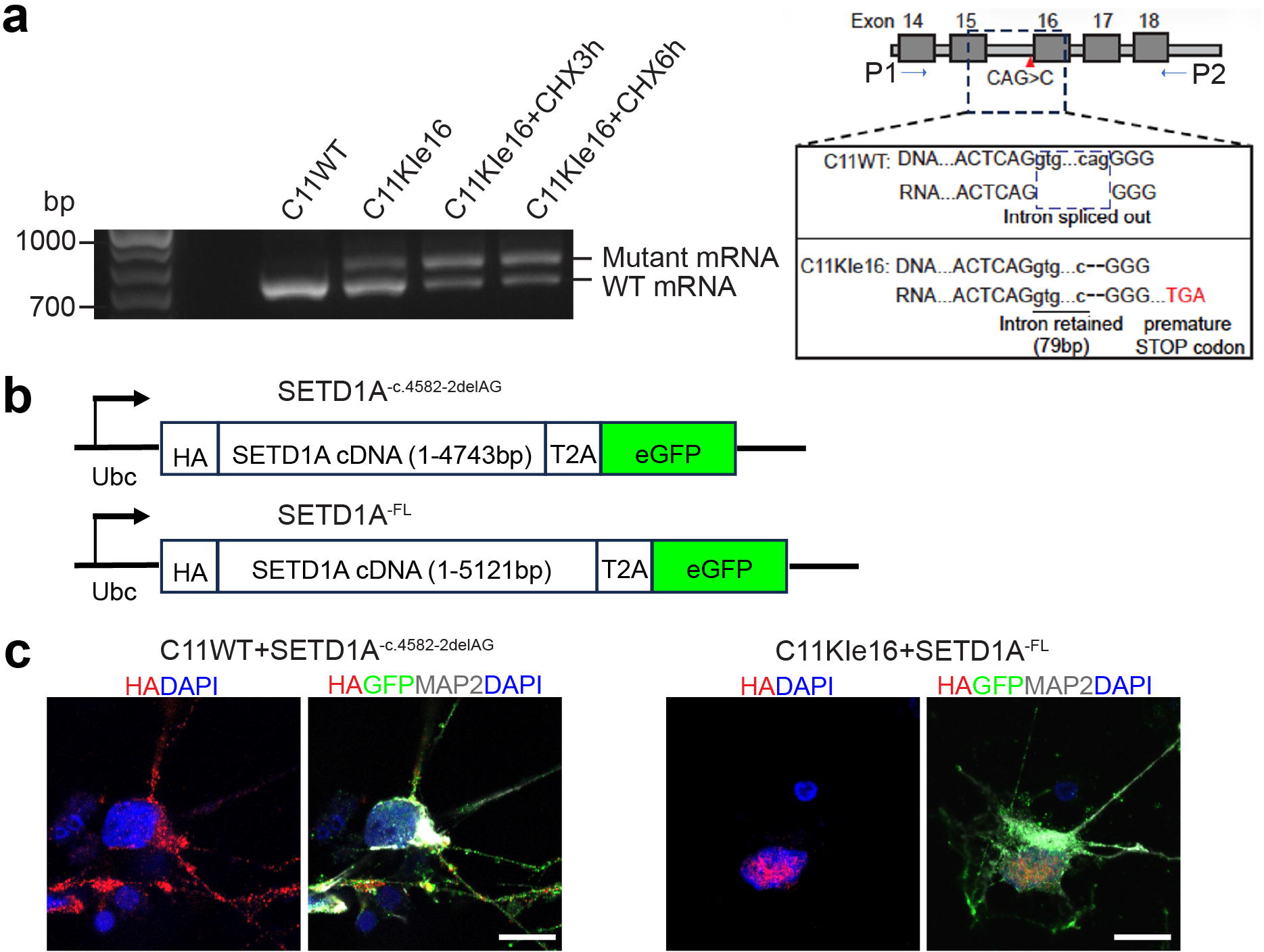
Expression of exon 16 patient-specific mutant SETD1A in DGC-like neurons. **a,** Endogenous exon 16 patient-specific mutant *SETD1A* transcripts existed in C11KIe16 neurons, and administration of cycloheximide (CHX), an inhibitor of nonsense-mediated decay, increased the amount of endogenous mutant transcripts in C11KIe16 neurons. P1, forward PCR primer; P2, reverse PCR primer**. b,** Schematic diagram of SETD1A constructs. **c,** HA immunostaining revealed aberrant cytoplasmic localization of exon 16 patient-specific mutant SETD1A overexpressed in DGC-like neurons (Scale bars: 10 μm).

**Extended Data Fig. 11.**
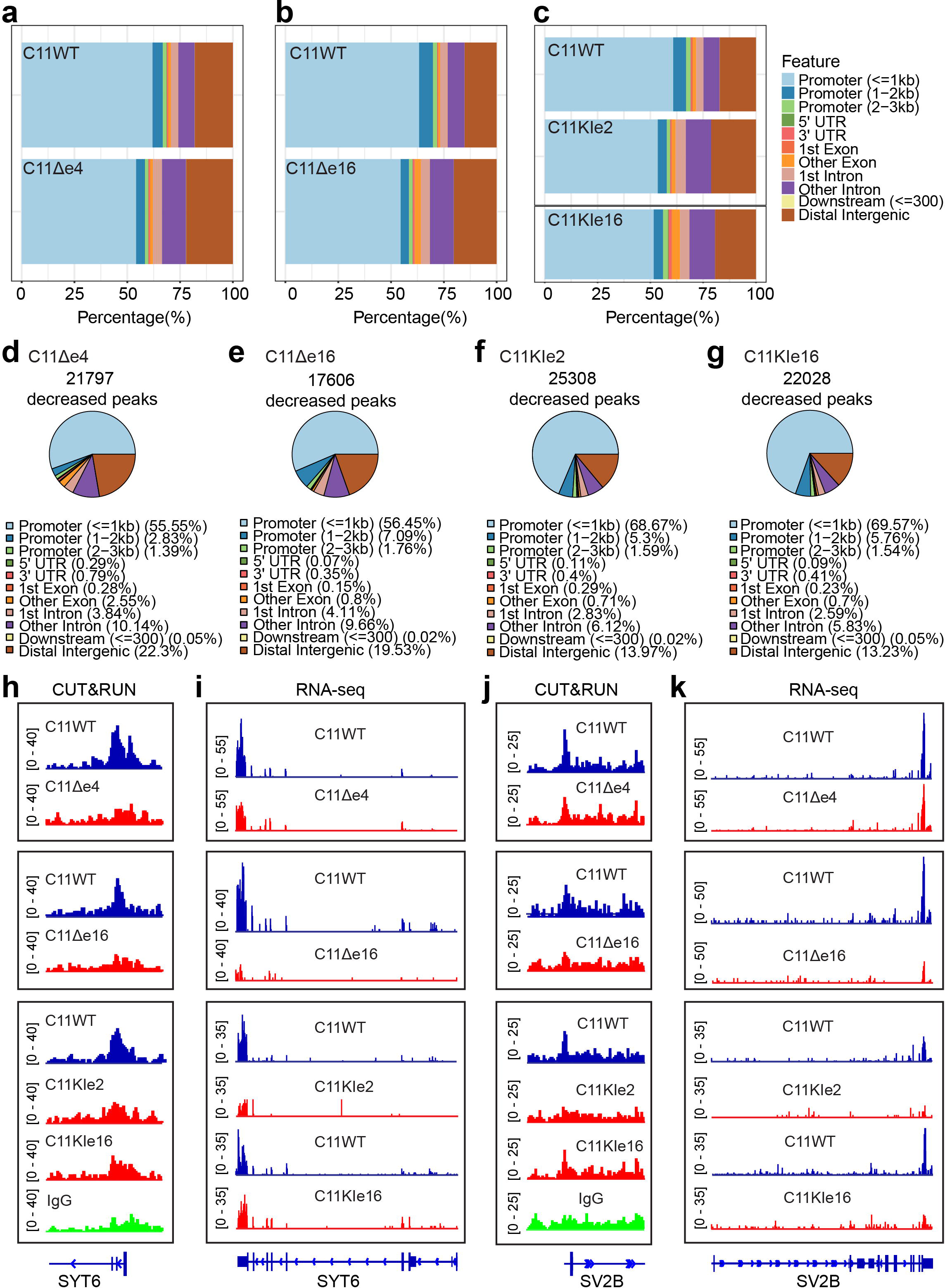
Genome-wide mapping of H3K4me3 in human DGC-like neurons with different *SETD1A* variants. **a-c,** Distribution of H3K4me3 peaks at the indicated genomic features in lines C11WT, C11Δe4, C11Δe16, C11KIe2, and C11KIe16. **d-g**, Distribution of reduced H3K4me3 peaks at the indicated genomic features in lines C11Δe4, C11Δe16, C11KIe2, and C11KIe16. **h, j,** Coverage plots from each line showing representative examples of H3K4me3 signals at the promoters of genes (*SYT6* and *SV2B*). **i, k,** RNA-seq tracks from each line showing the coverage of reads of genes (*SYT6* and *SV2B*).

**Extended Data Fig. 12.**
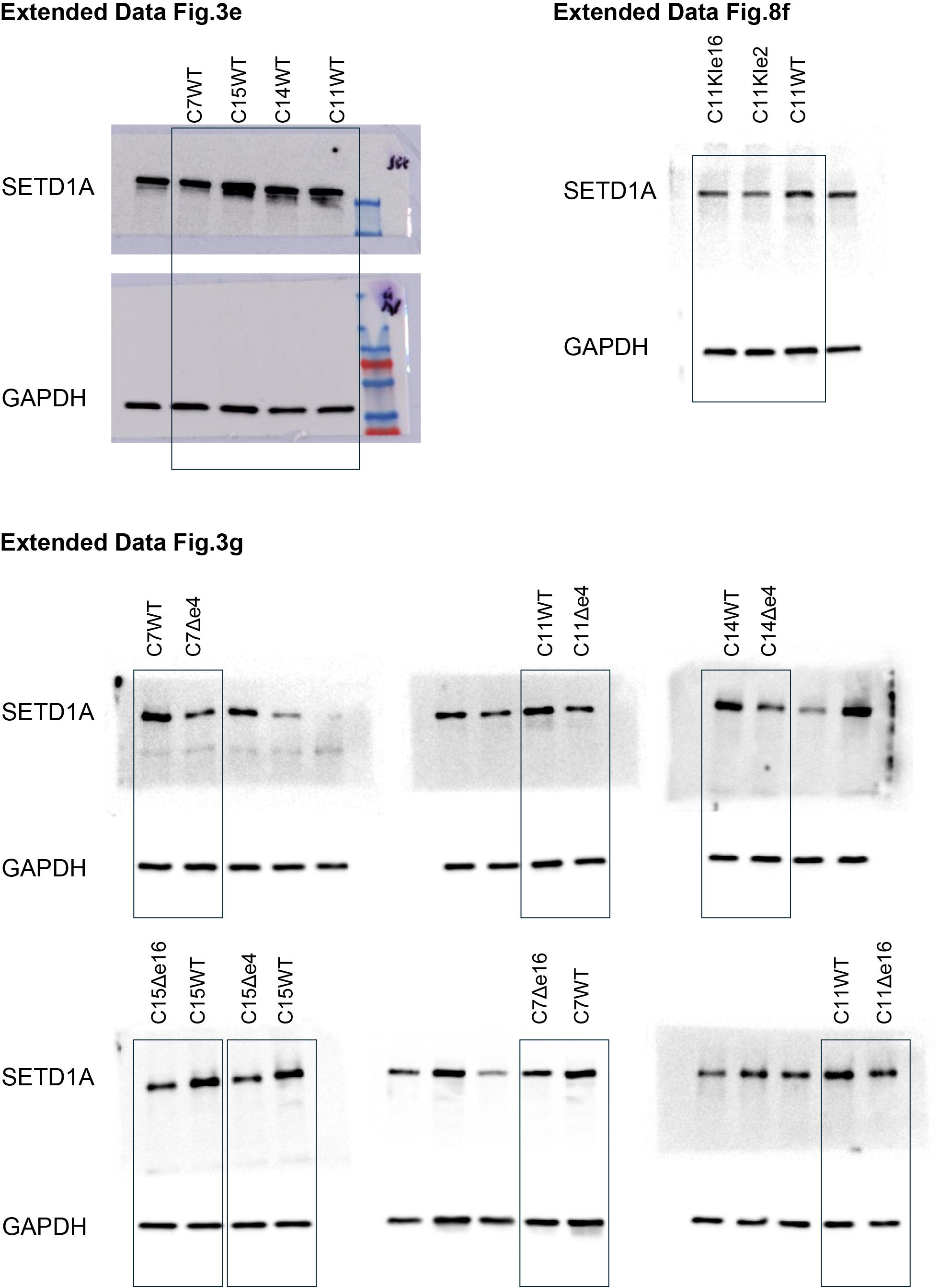
Whole gel images for Westernblot analysis.

## Extended Data Tables

**Extended Data Table 1. General information on cell lines, antibodies, and primer sequences. Extended Data Table 2. List of differentially expressed genes from RNA-seq data.**

**Extended Data Table 3. Decreased H3K4me3 peaks in different mutant lines with the same genetic background.**

**Extended Data Table 4. List of raw reads from RNA-seq data.**

